# An RNAi screen to identify proteins required for cohesion rejuvenation during meiotic prophase in Drosophila oocytes

**DOI:** 10.1101/2023.12.15.571853

**Authors:** Muhammad A. Haseeb, Alana C. Bernys, Erin E. Dickert, Sharon E. Bickel

## Abstract

Accurate chromosome segregation during meiosis requires maintenance of sister chromatid cohesion, initially established during premeiotic S phase. In human oocytes, DNA replication and cohesion establishment occur decades before chromosome segregation and deterioration of meiotic cohesion is one factor that leads to increased segregation errors as women age. Our previous work led us to propose that a cohesion rejuvenation program operates to establish new cohesive linkages during meiotic prophase in Drosophila oocytes and depends on the cohesin loader Nipped-B and the cohesion establishment factor Eco. In support of this model, we recently demonstrated that chromosome-associated cohesin turns over extensively during meiotic prophase and failure to load cohesin onto chromosomes after premeiotic S phase results in arm cohesion defects in Drosophila oocytes. To identify proteins required for prophase cohesion rejuvenation but not S phase establishment, we conducted a Gal4-UAS inducible RNAi screen that utilized two distinct germline drivers. Using this strategy, we identified 29 gene products for which hairpin expression during meiotic prophase, but not premeiotic S phase, significantly increased segregation errors. Prophase knockdown of Brahma or Pumilio, two positives with functional links to the cohesin loader, caused a significant elevation in the missegregation of recombinant homologs, a phenotype consistent with premature loss of arm cohesion. Moreover, fluorescence in situ hybridization confirmed that Brahma, Pumilio and Nipped-B are required during meiotic prophase for maintenance of arm cohesion. Our data support the model that Brahma and Pumilio regulate Nipped-B dependent cohesin loading during rejuvenation. Future analyses will better define the mechanism(s) that govern meiotic cohesion rejuvenation and whether additional prophase-specific positives function in this process.

## Introduction

In both mitotic and meiotic cells, accurate chromosome segregation requires that sister chromatids remain physically associated from the time of their synthesis (S phase) until they segregate to opposite poles (MCNICOLL *et al*. 2013; MARSTON 2014; MORALES AND LOSADA 2018; ISHIGURO 2019). In addition, during meiosis, cohesion between the arms of sister chromatids also provides an evolutionarily conserved mechanism to keep recombinant homologs associated until anaphase I (BUONOMO *et al*. 2000; BICKEL *et al*. 2002; HODGES *et al*. 2005). Within the cohesin complex, which mediates sister chromatid cohesion, association of an α-kleisin subunit with the Smc1/Smc3 heterodimer results in formation of a ring. Opening of the ring and topological entrapment of DNA by cohesin requires the cohesin loader (Scc2/Scc4 in yeast, NIPBL/MAU2 in mammals) (ALONSO-GIL AND LOSADA 2023). During DNA replication, formation of stable cohesive linkages depends on the acetyltransferase (Eco1 in yeast, ESCO1/2 in mammals) which keeps the ring stably closed by acetylating two conserved lysines within the Smc3 head (PETERS AND NISHIYAMA 2012; RANKIN AND DAWSON 2016).

In metazoans, one challenge oocytes face is that the formation of cohesive linkages during premeiotic S phase can occur days to decades prior to chromosome segregation, depending on the organism. Therefore, proper chromosome segregation in oocytes demands that a sufficient number of the original cohesive linkages remain intact or be replaced during the long prophase I arrest. Loss of cohesion in aging oocytes has been observed in multiple organisms (CHIANG *et al*. 2012; GREANEY *et al*. 2018; WARTOSCH *et al*. 2021; CHARALAMBOUS *et al*. 2023) and cohesin turnover on meiotic chromosomes has not been detected in mouse oocytes (REVENKOVA *et al*. 2010; TACHIBANA-KONWALSKI *et al*. 2010; BURKHARDT *et al*. 2016). These observations have led to the model that gradual deterioration of the original cohesive linkages in human oocytes contributes to increased segregation errors in the oocytes of older women, a phenomenon known as the maternal age effect.

Drosophila provides a powerful genetic system to dissect the mechanisms that influence cohesion maintenance in oocytes. Because the female germline matα-Gal4-VP16 driver is not expressed until after completion of premeiotic S phase (WENG *et al*. 2014), one can use this driver to ask whether knockdown (KD) of a gene product exclusively during meiotic prophase disrupts the *maintenance* of cohesion in Drosophila oocytes. Using this strategy, we previously demonstrated that knockdown of individual cohesin subunits, the Drosophila cohesin loader Nipped-B or the cohesion establishment factor Eco *after* premeiotic S phase results in phenotypes consistent with premature loss of cohesion (WENG *et al*. 2014). Based on these findings, we proposed that a cohesion rejuvenation program operates in Drosophila oocytes during meiotic prophase to establish new cohesive linkages that are required to maintain cohesion (WENG *et al*. 2014). In support of this hypothesis, we have recently reported that chromatin-associated cohesin turns over extensively during meiotic prophase in Drosophila oocytes. (HASEEB *et al*. 2023). Moreover, failure to load cohesin onto oocyte chromosomes during meiotic prophase leads to premature loss of arm cohesion (HASEEB *et al*. 2023). These data provide evidence that de novo formation of cohesive linkages occurs after S phase in Drosophila oocytes and is required to maintain the association of sister chromatids and support accurate chromosome segregation during the meiotic divisions.

Understanding the mechanism(s) underlying cohesion rejuvenation requires identification of the proteins involved, particularly those that may be unique to this process. Given that cohesion rejuvenation in the Drosophila oocyte occurs in the absence of global DNA replication, it likely differs mechanistically from the formation of stable cohesive linkages during S phase. In addition, we have previously shown that cohesion rejuvenation in prophase oocytes occurs in the absence of double-strand breaks (WENG *et al*. 2014) distinguishing it from the pathway that operates during G2 in mitotically dividing yeast cells subjected to DNA damage (STROM *et al*. 2004; STROM *et al*. 2007; UNAL *et al*. 2007).

With the goal of identifying proteins that are required for prophase rejuvenation but not S phase establishment, we designed a Gal4/UAS RNAi screen to quantify and compare chromosome segregation errors (NDJ) in control oocytes (hairpin but no driver), prophase KD oocytes (matα-Gal4-VP16→ hairpin) and S phase KD oocytes (nanos-Gal4-VP16→ hairpin) (see **Figs 1 & 2**). We reasoned that a hairpin targeting a protein *specific* for cohesion rejuvenation should cause a significant increase in NDJ when expressed during meiotic prophase but not premeiotic S phase.

**Figure 1.**
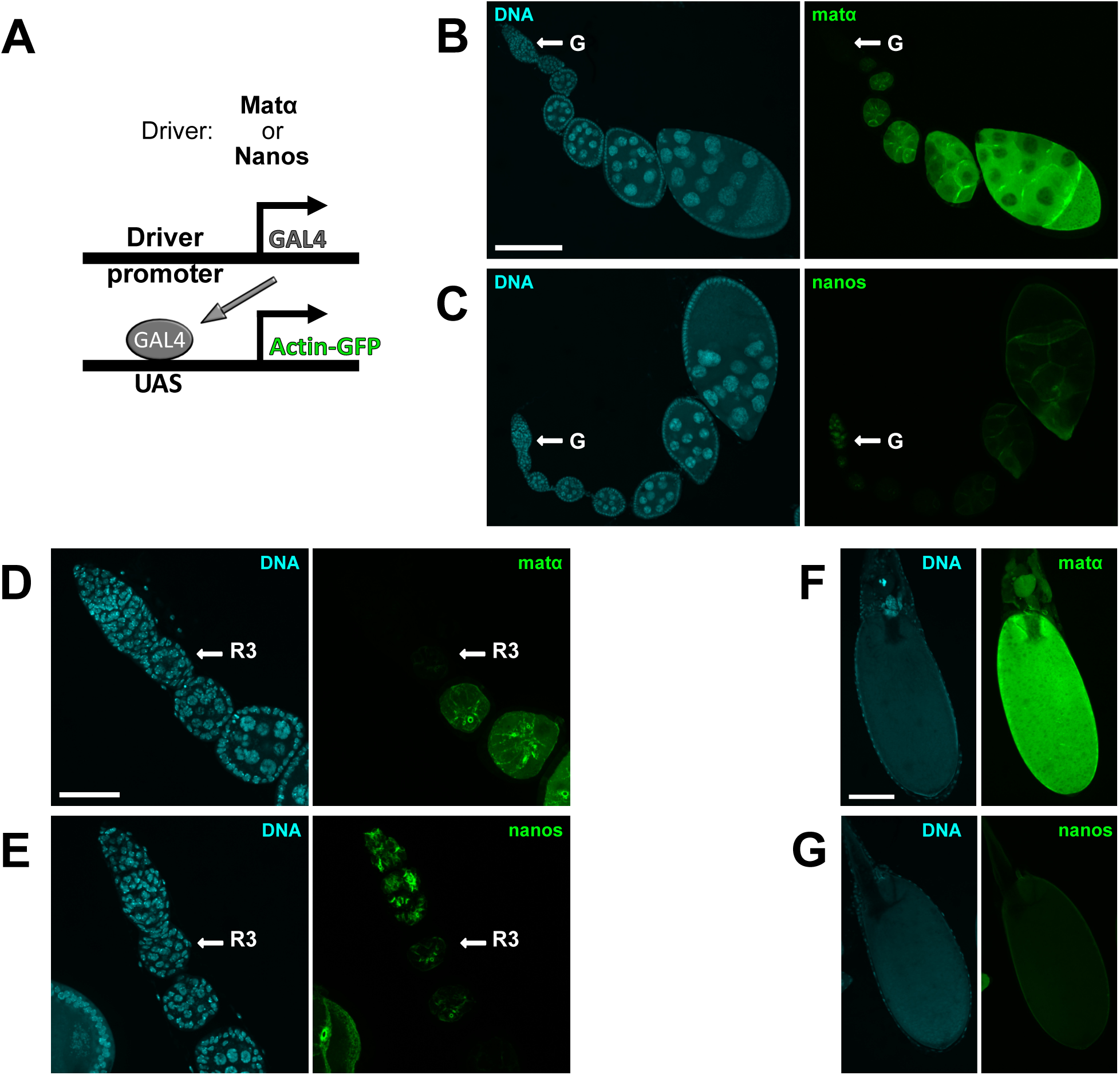
Comparison of matα-Gal4 and nanos-Gal4 expression patterns and relative strengths within Drosophila ovarioles. **A.** Schematic illustrates Gal4/UAS method utilized to express GFP-tagged actin in Drosophila ovaries using two different germline-specific drivers, matα-Gal4-VP16 or nanos-Gal4-VP16. **B-C**. Drosophila ovarioles are shown for which UASp-actin-GFP expression (green) is induced by the matα or the nanos driver. DNA is shown in blue. Arrows indicate the germarium (G) at the anterior of each ovariole. Scale bar, 120µm. To allow comparison of relative driver strengths, images for both drivers were captured and processed identically in this and subsequent panels. All images are maximum intensity projections of confocal Z series. **D-E.** Higher magnification images highlight the different expression patterns of the two drivers within the germarium. Region 3 (R3) of the germarium is labeled. Scale bar, 40µm. Note that nanos-Gal4 induced expression is visible within several germ-line cysts of the germarium, including the stage at which meiotic DNA replication occurs. In contrast, the matα-Gal4 driver does not turn on until R3 or Stage 2, approximately two days after premeiotic S phase. **F-G.** In mature oocytes (stages 13-14), actin-GFP signal resulting from the matα driver is much stronger than that induced by the nanos driver. Scale bar, 120µm.

**Figure 2.**
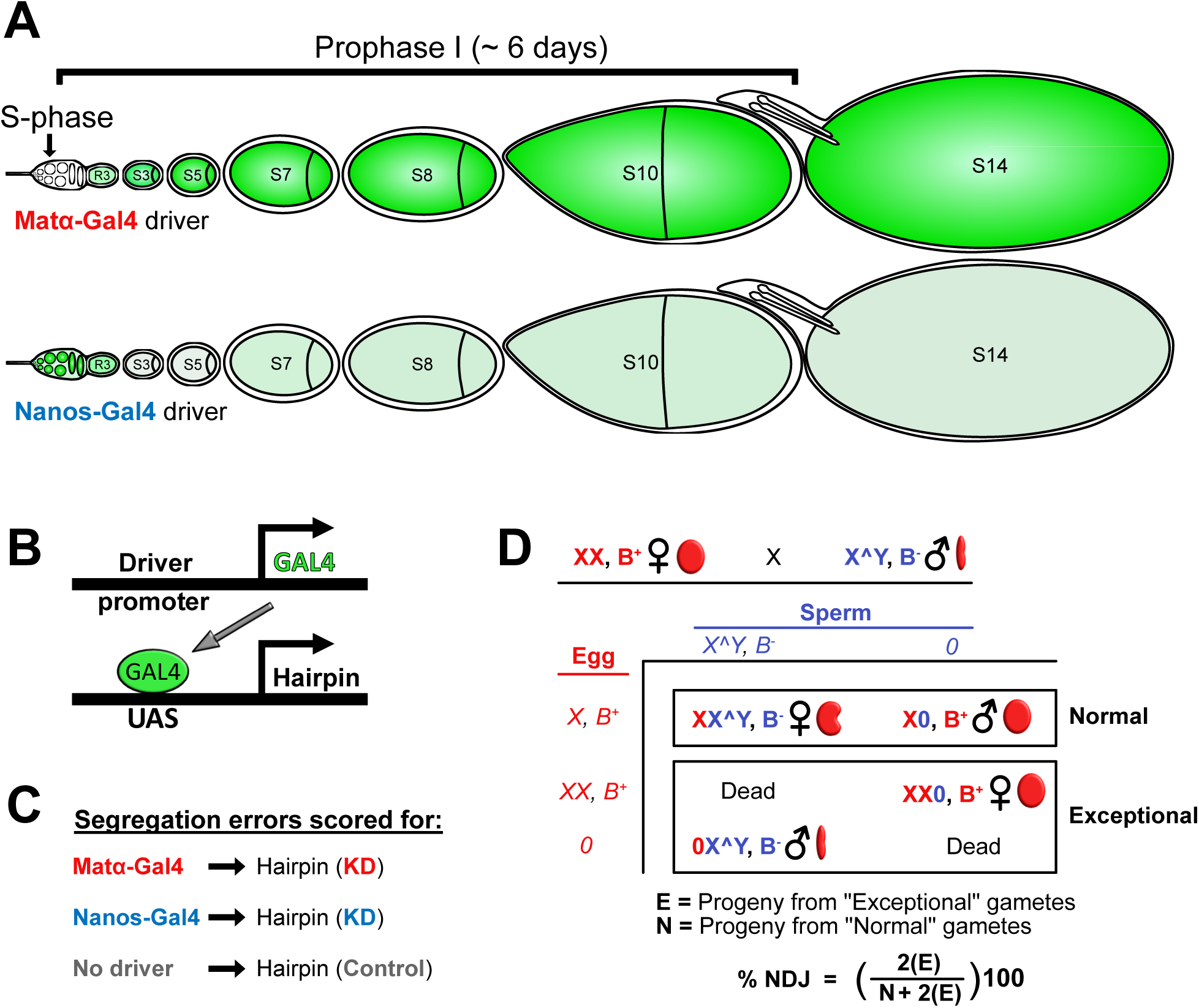
Experimental strategy to identify proteins required for cohesion rejuvenation during meiotic prophase. **A.** Cartoon representation of the Drosophila ovariole uses green shading to depict the expression patterns and relative strengths of the matα-Gal4 and nanos-Gal4 drivers. Upon completion of premeiotic S phase in the germarium, germline cysts enter meiotic prophase. Note that not all stages are present in a single ovariole at any given timepoint. **B.** UAS-Gal4 strategy utilizes the matα-Gal4 or nanos-Gal4 driver to express a specific hairpin in the female germline. **C.** Chromosome segregation errors in knockdown (KD) oocytes are quantified and compared to control oocytes containing the hairpin transgene but no driver. **D.** In the *X*-chromosome nondisjunction (NDJ) assay, KD or control females are crossed to males with an attached *X^Y* chromosome containing the dominant eye-shape marker, Bar. Based on their sex and eye shape, progeny arising from accurate *X* chromosome segregation (Normal, N) can be distinguished from those resulting from aberrant segregation (Exceptional, E). Because only half of the exceptional gametes result in viable progeny, the value of E is doubled when calculating total % NDJ.

Here we report the results of our screen, including identification of 29 positives for which knockdown with the matα but not the nanos driver significantly increases meiotic chromosome segregation errors. Functional links to the cohesin loader have been reported for two prophase-specific positives, Brahma (Brm) and Pumilio (Pum). Knockdown of either protein during meiotic prophase caused a significant elevation in the frequency at which recombinant homologs missegregate, a phenotype consistent with premature loss of arm cohesion. Furthermore, direct analysis of the state of cohesion using fluorescence in situ hybridization (FISH) indicated that matα-induced KD of Brm or Pum causes a significant increase in oocytes with premature loss of arm cohesion. These results validate the general strategy of our screen and suggest that further analysis of additional positives will provide insight into the mechanism(s) underlying cohesion rejuvenation during meiotic prophase.

## Materials and Methods

### Fly stocks and crosses

All fly stocks and crosses were maintained on standard cornmeal-molasses food at 25°C in a humidified incubator. **Table S1** provides genotypes and information for general stocks utilized in the screen and specific hairpin stocks used for follow-up experiments. **Table S2** provides stock information for each of the hairpin stocks tested for NDJ in the primary screen.

### *X*-chromosome NDJ assay

Figure S1 shows representative crosses performed in parallel for each hairpin tested. Males carrying a UAS-hairpin transgene were crossed to *y w; + ; mtrm^KG^ matα-Gal4-VP16 /TM3* (W-110), *nanos-Gal4-VP16; + ; mtrm^KG^/TM3 (*T-764) *or y w; + ; mtrm^KG^/TM3* virgins (W-109). For the nondisjunction (NDJ) assay, non-balancer virgin female progeny from the above crosses were mated to *X^Y, v f B* (C-200) males. For each of the three genotypes tested for each hairpin (**Fig 2C**, **Fig S1**), 10 vials of the NDJ cross were started (8 virgins x 4 males) and oocyte segregation errors were measured by scoring the progeny through day 18 (**Fig 2D**). P values were calculated using the method described in ZENG *et al*. (2010).

### *X*-chromosome recombinational history and crossover frequency assays

To determine whether matα-induced KD of Brm or Pum increased missegregation of recombinant homologs, we utilized a genetic “recombinational history" assay that allows us to deduce whether an individual Diplo-*X* female arising from a NDJ event carries two homologs or two sister chromatids and whether one or both *X* chromosomes underwent a crossover before missegregation (SUBRAMANIAN AND BICKEL 2008; WENG *et al*. 2014; PERKINS *et al*. 2016). We created Brm and Pum hairpin stocks (I-563 & I-576, **Table S1**) that contained an *X* chromosome marked with *y*.

Virgins from these stocks were crossed to *y sc cv v f car/B[S]Y; + ; mtrm^KG^/TM3* (M-835) and *y sc cv v f car/B[S]Y; + ; matα mtrm^KG^/TM3* (M-834) males. The resulting *y/y sc cv v f car* ; *mtrm^KG^/TRiP hairpin* (**control**) and *y/y sc cv v f car* ; *matα mtrm^KG^/TRiP hairpin* (**KD**) virgins were crossed to *X^Y, v f B* males (C-200) and NDJ scored daily from day 10 through day 18. Diplo-*X* progeny were collected each day, phenotyped for *sc, cv, f,* and *car,* and each female mated to two *y w* males (A-062). By scoring her male progeny for *sc, cv, f,* and *car a*nd considering the phenotype of the Diplo-*X* female, we were able to deduce the genotype of the two *X* chromosomes she inherited, whether they were sisters or homologs (based on *car*) and whether either were recombinant (**Fig S2**). The frequency at which recombinant chromosomes missegregated was calculated by dividing the number of Diplo-*X* progeny that inherited at least one recombinant chromosome by the total number of progeny in the NDJ test and multiplying by 1000 to facilitate comparisons. P values were calculated using a two-tailed Fisher’s exact test (GraphPad).

One limitation of this assay is that it underestimates the number of recombinant bivalents that underwent missegregation because only two of the four chromatids can be genotyped. In addition, double crossovers in the large interval between *cv* and *f* will be invisible to us. Finally, although the proximity of *car* to pericentric heterochromatin (3.5cM) makes crossovers unlikely, a small number may still occur.

To determine whether knockdown of Brm or Pum altered the frequency and/or distribution of *X* chromosome crossovers, we crossed *y/y sc cv v f car* ; *mtrm^KG^/TRiP hairpin* (**control**) and *y/y sc cv v f car* ; *matα mtrm^KG^/TRiP hairpin* (**KD**) virgins to *y w* males (A-062). Male progeny were scored for *sc*, *cv*, *f* and *car* and map distance was calculated for each interval. A two-tailed Fisher’s exact test (GraphPad) was used to calculate significance.

### Whole mount ovary preparation for GFP-tagged actin reporter expression

To characterize the expression patterns and relative strengths of the two germline Gal4 drivers used in this study, we analyzed ovaries from flies that expressed GFP-tagged actin under the control of the matα or nanos Gal4 driver. *y w; + ; mtrm^KG^ matα-Gal4-VP16 /TM3* (W-110) or *nanos-Gal4-VP16; + ; mtrm^KG^/TM3 (*T-764) males were mated to *y w; P{UASp-Act5C.T:GFP}2; +* (A-201) virgins. Non-balancer young female progeny were held in food vials with males and dry yeast for two days before dissection in a shallow dish containing 1X PBS. The anterior region of each ovary was gently splayed open and ovaries fixed for five minutes at room temperature in 1X PBS containing 4% formaldehyde (Ted Pella, 18505). After three rinses in 1X PBS, ovaries were incubated in 1X PBS containing 2.0µg/ml Hoechst 33342 (Molecular Probes H3570) with gentle shaking for 30 minutes. Following three rinses and a 15 minute wash in 1X PBS, individual ovarioles were separated using tungsten needles and transferred to poly-L-lysine coated 18mm #1.5 coverslips. 25 µl of SlowFade Diamond Antifade (Molecular Probes S36967) was used for mounting and edges of the coverslips were sealed with nail polish before imaging.

### Fluorescence *in situ* hybridization (FISH)

We utilized FISH to quantify cohesion defects in KD and control oocytes containing Brm, Pum or Nipped-B hairpin transgenes. To generate each pair of samples (KD and control), *y sc v; + ;P{TRiP Brm^V20^}attP2* (H-209), *y sc v sev; + ;P{TRiP Pum^V20^}attP2* (H-211), or *y sc v; P{TRiP Nipped-B^V22^}attP40; +* (H-063) virgins were crossed to *w; + ; P{matα-GAL4-VP16}V37* (T-273) or *y w ; + ; +* (A-062) males. Note that oocytes used for all FISH experiments were wildtype for *mtrm*. Young female progeny were held in food vials with males and yeast for three days before ovaries were dissected. After fixation, stage 13-14 oocytes were processed as previously described (PERKINS *et al*. 2016; PERKINS AND BICKEL 2017). Fixation, pre-denaturation, hybridization, washes and mounting were performed as reported in HASEEB *et al*. (2023) except that ovaries were fixed for six minutes instead of four minutes. To monitor arm cohesion, we utilized an Alexa 647-labeled Oligopaint probe (OPP122 from the Joyce Lab, University of Pennsylvania) comprised of a mixture of 80-base oligonucleotides that hybridize across a 100kb distal region on *X* chromosome. Cohesion within the pericentric heterochromatin was analyzed using a Cy3-conjugated probe (5′-Cy3-AGGGATCGTTAGCACTCGTAAT; Integrated DNA Technologies) that targets an 11Mb region of satellite DNA on the *X* chromosome (DERNBURG 2000). Arm and pericentric probes were used at final concentrations of 0.50pmol/μl and 1ng/μl, respectively.

Following image acquisition, cohesion defects were scored, blind to genotype, as detailed in HASEEB *et al*. (2023). A two-tailed Fisher’s exact test (Graphpad) was used to determine the statistical significance for differences between KD and control.

### Image acquisition and analysis

All images were acquired using an Andor spinning disk confocal on a Nikon Eclipse Ti inverted microscope equipped with an ASI MS-2000 motorized piezo stage, a 50µm pinhole disk, and a Zyla 4.2-megapixel sCMOS camera. Nikon elements software (version 5.11.02 Build 1369) and up to four lasers (405, 488, 561 and 637 nm) were used for image acquisition. For Figure 1D & E, we utilized a Nikon CFI 40X Plan Fluor oil objective (N.A. 1.3) to acquire a specified region of interest. For Figure 1B & C and 1F & G, full-frame images were captured using a Nikon CFI 20X Plan Apo objective (N.A. 0.75). All FISH images were collected using a Nikon CFI 100X oil Plan Apo DIC objective (NA 1.45). 4X frame averaging was employed for all image acquisition. For Z-series, a step size of 1 µm, covering an 8 µm range was used for Figure 1, and a 0.1 µm step size over a 4 µm range was utilized for FISH imaging. Starting with the longest wavelength, an entire Z-stack was acquired with one laser before proceeding to the next channel. Images presented in Figure 1 were acquired and processed identically for both matα-Gal4 and nanos-Gal4 driven UASp-actin-GFP, including the number of optical sections included in the projections.

## Results and Discusion

### Screen rationale and design

To screen for proteins required for cohesion maintenance during meiotic prophase, we utilized two different germ-line specific Gal4-VP16 drivers to induce expression of RNAi hairpins at different times during Drosophila oogenesis. Fig 1 compares the relative strengths and expression patterns of the matα-Gal4-VP16 and nanos-Gal4-VP16 drivers (hereafter referred to as matα and nanos drivers). Expression of the matα driver is first detectable in germarial Region 3 (WENG *et al*. 2014) or stage 2 of the ovariole (Fig 1), approximately two days after completion of premeiotic S phase. Therefore, knockdown using this driver is restricted to meiotic prophase. In contrast, the nanos driver is expressed in multiple mitotic and meiotic cysts within the germarium, including the cells undergoing premeiotic S phase (Fig 1). Although nanos-Gal4 induced expression is also detectable during mid to late prophase, it is substantially weaker than that for the matα driver (Fig 1).

We utilized *mtrm^KG08051/+^* heterozygotes as a sensitized genetic background for our screen. Matrimony protein is required for accurate segregation of achiasmate bivalents in Drosophila oocytes. This achiasmate segregation system, which relies on pericentric heterochromatin mediated association of homologs (HAWLEY *et al*. 1992; KARPEN *et al*. 1996), also ensures proper segregation of crossover homologs that lose their physical connection due to premature loss of arm cohesion (**Fig S3).** Therefore, hairpin expression that causes premature loss of meiotic cohesion may not significantly elevate NDJ if oocytes are wild-type for *mtrm.* Because the achiasmate pathway is disabled in *mtrm^KG/+^* oocytes (HARRIS *et al*. 2003), our ability to identify gene products required for cohesion maintenance is enhanced in this genotype. In addition to its role in achiasmate segregation, Mtrm also promotes maintenance of sister chromatid cohesion in late prophase by binding and inhibiting Polo kinase (XIANG *et al*. 2007; BONNER *et al*. 2020; HASEEB *et al*. 2023). Although we have observed (using FISH) that arm cohesion defects are higher in *mtrm^KG/+^* oocytes than *mtrm^+^* oocytes, weak expression of a hairpin that targets the cohesin subunit Smc3 significantly increases cohesion defects in *mtrm^KG/+^* oocytes (HASEEB *et al*. 2023). Therefore, weak cohesion defects in KD oocytes will likely be amplified in *mtrm^KG/+^*oocytes, increasing the likelihood of elevated NDJ (**Fig S3**).

To generate a list of potential gene products to knock down in our screen, we searched Flybase for genes expressed in the female germline (oocyte and/or nurse cells). For each candidate on this list, we determined whether a Valium 20 or 22 insertion was available from the TRiP (Transgenic RNAi Project) collection (NI *et al*. 2011). These vectors provide robust hairpin expression and effective knockdown in the germline. We favored UAS-hairpins inserted at the attP2 site (chromosome *3*), but also tested a few attP40 insertions (chromosome *2*). For each candidate tested, we performed crosses (**Fig S1**) to generate control (no KD), matα-driven KD and nanos-driven KD oocytes and performed our NDJ assay in parallel for all three genotypes (Fig 2).

### Hairpins for which nanos-Gal4 induced knockdown causes a significant increase in meiotic segregation errors

Given that the cohesin complex is required for cohesion establishment during S phase, we utilized hairpins targeting the cohesin subunits Smc1 and Smc3 to first validate our approach. **Fig 3A** compares NDJ in oocytes in which one of these cohesin subunits was knocked down using either the nanos or matα driver. Relative to control oocytes, nanos-induced KD of either cohesin subunit resulted in a robust, statistically significant increase in NDJ, consistent with their essential role during S-phase cohesion establishment. In addition, as we have reported previously (WENG *et al*. 2014), matα-induced KD of Smc1 or Smc3 also caused a significant elevation in meiotic NDJ compared to control, consistent with their requirement for cohesion rejuvenation during meiotic prophase. Notably, for both Smc1 and Smc3, NDJ was significantly higher for nanos KD than matα KD (see **Table S3** for P values). We consider the above phenotypes to be a useful reference indicative of gene products that are required for both S phase establishment and prophase rejuvenation in Drosophila oocytes.

**Figure 3.**
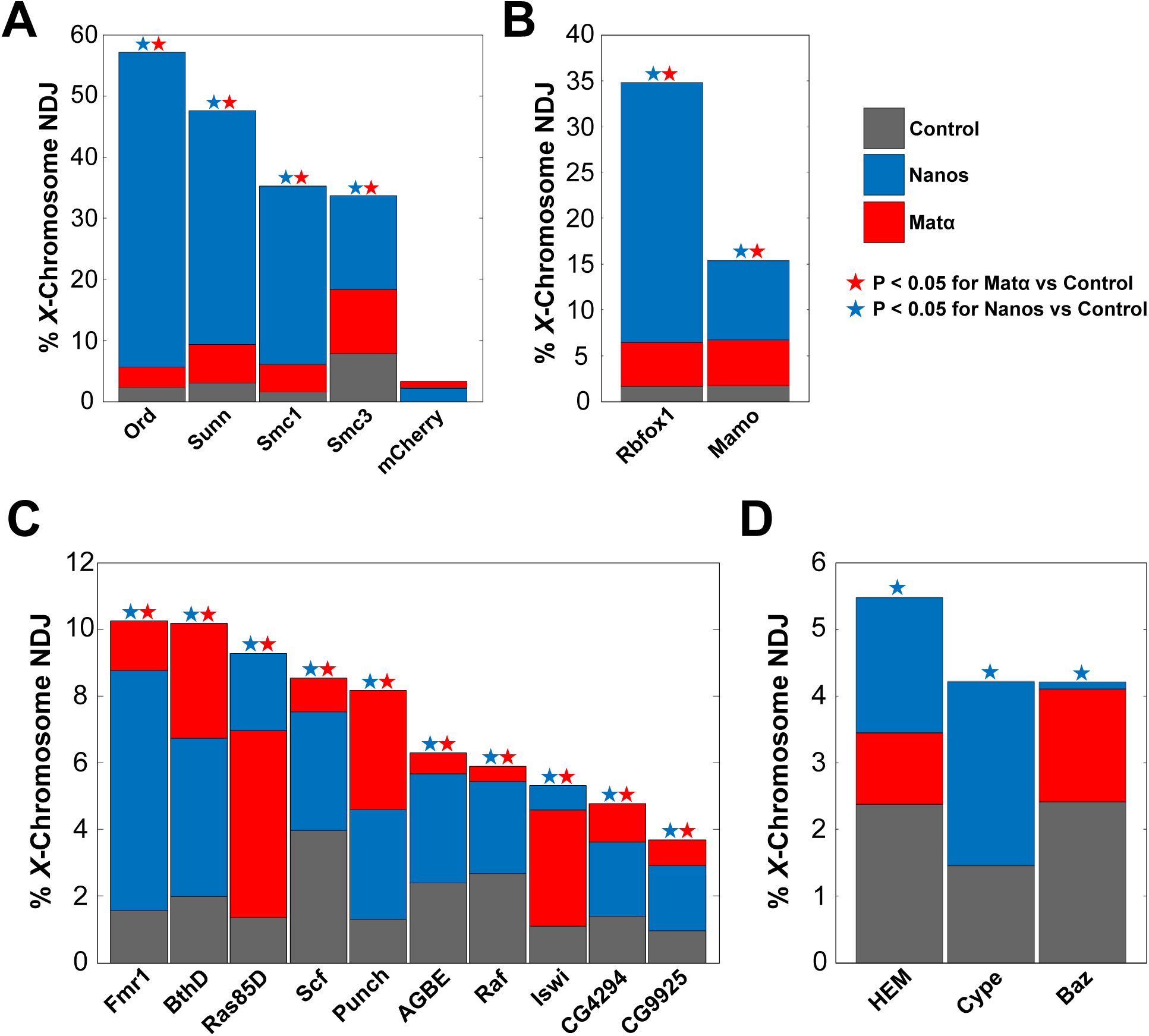
Gene products for which nanos-induced knockdown (KD) causes a significant elevation in NDJ. Overlaid bars are used to graph *X*-chromosome NDJ in no driver → control (gray), nanos-Gal4 → KD (blue), and matα-Gal4 → KD (red) oocytes. Each bar starts at zero on the y-axis and its top height corresponds to the NDJ value for that color. A colored star indicates a significant difference (P < 0.05) between the NDJ in control oocytes and those with the indicated driver (blue star, nanos; red star, matα). **A.** NDJ increases significantly when either the nanos or matα driver is used to knock down Drosophila proteins known to be required for meiotic cohesion. However, NDJ is significantly higher with S phase KD than prophase KD (Table S3). Expression of an mCherry hairpin serves as a negative control. **B.** Knockdown of Rbfox1 or Mamo using either driver causes a significant increase in NDJ but segregation errors are significantly greater for nanos-induced KD than for matα-induced KD (Table S3). **C.** Knockdown using either driver elicits a significant increase in NDJ that is comparable for the two drivers (Table S4). **D.** Gene products for which knockdown using the nanos driver but not the matα driver significantly elevates NDJ (Table S5).

As a negative control, we expressed a Valium 20 hairpin targeting mCherry in flies that lack a mCherry-encoding transgene (**Fig 3A**). Compared to the no driver control, meiotic NDJ was not significantly elevated with either the nanos or the matα drivers in *mtrm^KG/+^* heterozygotes containing the mCherry hairpin (**Fig 3A, Table S3**). These results strengthen our confidence that candidates uncovered in our screen are not false positives.

As part of our validation strategy, we also investigated two other proteins required for meiotic cohesion in Drosophila for which Valium 20-22 hairpin stocks are available. Null mutations in the *orientation disruptor* (*ord*) gene cause segregation defects consistent with complete loss of meiotic cohesion in both oocytes and spermatocytes (BICKEL *et al*. 1997). Although the molecular function of Ord is not fully understood, Ord protein localizes to both the arms and centromeres of meiotic chromosomes in Drosophila oocytes (KHETANI AND BICKEL 2007), is required for chiasma maintenance (BICKEL *et al*. 2002) and for localization of Smc1 and Smc3 to oocyte centromeres (WEBBER *et al*. 2004). Mutations in *sisters unbound* (*sunn*) also disrupt meiotic cohesion in both oocytes and spermatocytes (KRISHNAN *et al*. 2014) and Sunn protein has been proposed to function within one of the two meiosis-specific cohesin complexes in Drosophila (KRISHNAN *et al*. 2014; GYURICZA *et al*. 2016). Like Smc1 or Smc3 KD oocytes, meiotic NDJ is significantly elevated when Ord or Sunn is knocked down using either the nanos or the matα driver **(Fig 3A)**. Moreover, chromosome segregation errors are significantly more prevalent following S phase KD than prophase KD (see **Table S3** for P values). The data obtained with the nanos driver were not unexpected and indicate that like Smc1 and Smc3, both Ord and Sunn are essential for cohesion establishment during premeiotic S phase in Drosophila oocytes. In addition, our results using the matα driver suggest that Ord and Sunn proteins are also required during meiotic prophase for cohesion rejuvenation.

Interestingly, our screen uncovered two gene products for which knockdown resulted in phenotypes similar to those of the cohesion proteins tested above (**Fig 3B**, **Table S3**). Compared to control oocytes, nanos and matα drivers both caused a significant increase in meiotic NDJ in Rbfox1 (RNA-binding Fox protein 1) and Mamo (Maternal gene required for meiosis) KD oocytes. Although neither protein has been implicated in cohesion regulation, Mamo is required for normal chromatin structure in Drosophila oocytes (MUKAI *et al*. 2007; HIRA *et al*. 2013) and Rbfox1 regulates translation of Pumilio, one of the prophase-specific positives we discuss below (CARREIRA-ROSARIO *et al*. 2016). Our results raise the possibility that these proteins play a role in cohesion establishment during premeiotic S phase as well as cohesion rejuvenation during meiotic prophase.

For ten positives, NDJ was significantly elevated for both nanos and matα drivers (**Fig 3C**); however, for all but Punch, segregation errors did not differ significantly between the two drivers (**Table S4)**. In addition, nanos-induced KD of these proteins resulted in NDJ that was considerably lower than that observed for Smc1 or Smc3 KD (approximately 4-10 fold lower). The positives presented in **Fig 3C** have diverse functions and include two uncharacterized gene products (CG4294 and CG9925). We cannot rule out the possibility that activity of these proteins during premeiotic S phase is necessary for accurate chromosome segregation. However, although nanos-driven UASp-actin-GFP expression during prophase is considerably weaker than that for the matα driver, signal is visible after exit from the germarium (**Fig 1E**) and increases slightly in late prophase (**Fig 1C**). Therefore, weak knockdown of these proteins during meiotic prophase (not S phase) may result in the increased NDJ observed with the nanos driver.

Our screen also uncovered three proteins for which knockdown with the nanos but not the matα driver caused a relatively small, but significant increase in *X*-chromosome NDJ compared to no driver controls (Fig 3D, **Table S5**). Because the much stronger matα driver did not significantly increase NDJ compared to control oocytes, the phenotype obtained with the nanos driver is consistent with an essential role during premeiotic S phase. However, we cannot rule out an early prophase function or the possibility that knockdown in the mitotic cysts of the germarium is responsible for the increased meiotic NDJ we observe with the nanos driver.

### Prophase-specific positives

Of the 63 hairpin targets that we tested, knockdown of 29 proteins resulted in a significant increase in NDJ with the matα driver but not the nanos driver (Fig 4, **Table S6**). This prophase-specific phenotype is what we would expect for proteins that are required for cohesion rejuvenation during meiotic prophase and not for cohesion establishment during premeiotic S phase. Although matα-induced KD uncovered several negatives (**Table S7**), the high percentage of prophase-specific positives was unexpected (Fig 4, **Table S6**). However, because accurate chromosome segregation in oocytes depends on several pathways in addition to cohesion maintenance, it is unlikely that increased NDJ for all these hits arises because of premature loss of meiotic cohesion. As shown in **Fig 4C** and **Table S8**, the prophase-specific positives function in a diverse array of cellular pathways/mechanisms and include nine uncharacterized gene products. Most of these rejuvenation candidates exhibit less than 10% NDJ when knocked down using the matα driver, with only seven positives that exceed that value **(Fig 4A)**. However, this relatively low level of NDJ may still reflect a *bona fide* role in cohesion rejuvenation given that it is comparable to what we observe for matα-induced KD of three of the four cohesion proteins we tested (**Fig 3A)**. Although four hairpins (Hang, eIF5, Dhc64C, Hip14) resulted in a significant NDJ increase for matα-but not nanos-induced KD, we did not graph these as prophase-specific positives in **Fig 4A& B** because nanos→ KD oocytes were sterile or nearly sterile (**Table S6**). Such a phenotype is consistent with an essential role during premeiotic S phase.

**Figure 4.**
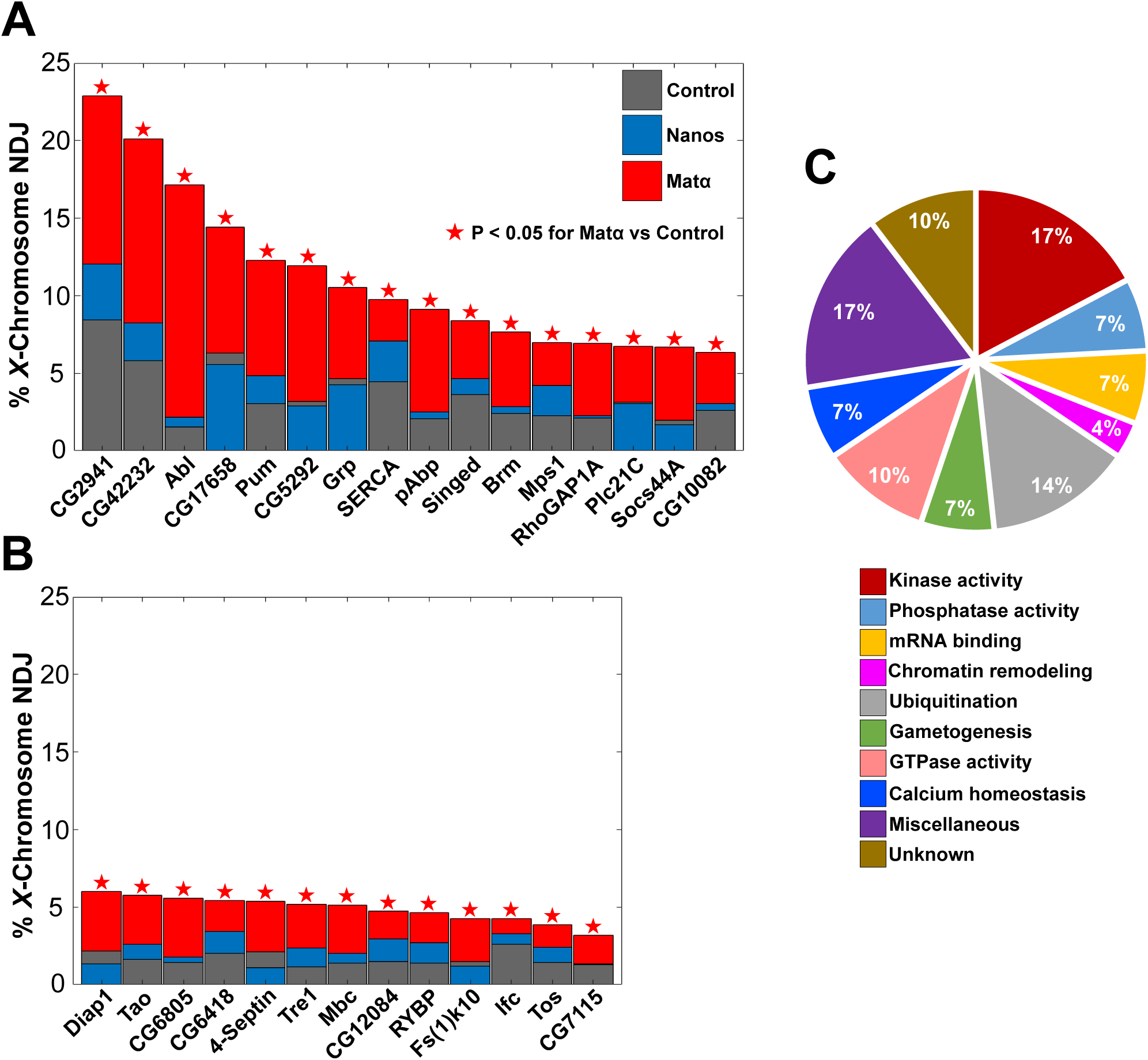
Proteins for which matα-induced but not nanos-induced knockdown causes meiotic NDJ to increase significantly. Overlaid bars are used in the same manner as in Fig 3. **A & B.** A significant increase in NDJ with the matα but not the nanos driver is consistent with a rejuvenation-specific role that is not required for cohesion establishment during oocyte DNA replication. **C.** Best-described cellular function for each of the prophase-specific positives is presented with a corresponding pie chart that indicates the relative percentage of positives in each category (also see Table S8).

For a subset of the positives shown in **Fig 4A-B**, we tested a second hairpin to potentially rule out off-target effects. For Abl (Ableson kinase), Brm (Brahma), Mps1 (Monopolar Spindle 1) and CG5292, NDJ arising with the second hairpin was consistent with our initial observations, significantly increased with the matα but not the nanos driver (**Table S6**). For CG10081 and CG6805, neither the matα nor the nanos driver caused a significant increase in NDJ with the second hairpin (**Table S6**). However, for these two uncharacterized genes, we cannot distinguish between an off-site target effect with the first hairpin or insufficient KD with the second hairpin to elicit a significant increase in NDJ. Interestingly, the 2^nd^ hairpin we utilized to knock down Pum (Pumilio) resulted in a significant NDJ elevation with the nanos driver as well as the matα driver, but NDJ was significantly lower for nanos-induced KD than for matα-induced KD (**Table S6**). As discussed above for the positives in **Fig 3C**, this phenotype could arise because of nanos driver activity in prophase and not reflect an essential role during premeiotic S phase.

### Brahma and Pumilio are required to maintain arm cohesion during meiotic prophase

Because several mechanisms govern accurate chromosome segregation, we set out to determine whether the NDJ we observed for a subset of prophase-specific positives occurs due to premature loss of cohesion. We selected Brahma (Brm) and Pumilio (Pum) for further analyses because each has published functional links with the cohesin loader (GERBER *et al*. 2006; MUNOZ *et al*. 2019; MUNOZ *et al*. 2020). Brm is the ATPase subunit of two chromatin remodeling complexes in Drosophila (CLAPIER AND CAIRNS 2009). In yeast, the remodeling complex that contains the Brm ortholog facilitates cohesin loading to nucleosome-free regions by recruiting the cohesin loader Scc2 (Nipped-B in Drosophila) (MUNOZ *et al*. 2019; MUNOZ *et al*. 2020). Pum belongs to the evolutionarily-conserved PUF family of sequence-specific RNA binding proteins that control protein abundance by regulating mRNA stability and/or translation (NISHANTH AND SIMON 2020). Interestingly, Pum has been shown to bind Nipped-B mRNA in Drosophila ovary extracts (GERBER *et al*. 2006).

Given that premature loss of arm cohesion leads to chiasma destabilization (BUONOMO *et al*. 2000; BICKEL *et al*. 2002; HODGES *et al*. 2005), we asked whether prophase KD of Brm or Pum increases missegregation of recombinant homologs (**Fig S2**). Using females that were heterozygous for recessive visible markers along the *X* chromosome, we repeated the NDJ assay and again observed a significant elevation in segregation errors in matα KD oocytes compared to their respective controls (**Fig 5A-B**). Following the NDJ assay, we performed an additional cross and used the *X* chromosome visible markers to genotype the male progeny for each Diplo-*X* female and determine whether either of the missegregating *X* chromosomes that she inherited were recombinant (**Fig S2**). The centromere-proximal marker *carnation* allowed us to determine whether the Diplo-*X* female inherited two homologs (*car ^+/-^,* MI error) or two sisters (*car ^+/+^*or *car ^-/-^*, MII error). We found that prophase KD of either Brm or Pum significantly increased the frequency at which Diplo-*X* females inherited two homologs (MI errors), at least one of which was recombinant (**Fig 5C-D**). MII errors (sisters) arising from a recombinant bivalent were not significantly different between KD and control oocytes (**Fig 5C-D**). Furthermore, the total map distance of the *X* chromosome was not significantly altered in KD oocytes (**Fig 5E-F**), indicating that missegregation of recombinant homologs is not elevated in Brm and Pum KD oocytes due to increased crossovers in these genotypes. Together, these data support the hypothesis that Brm and Pum are required for accurate chromosome segregation in Drosophila oocytes because they promote cohesion rejuvenation during meiotic prophase.

**Figure 5.**
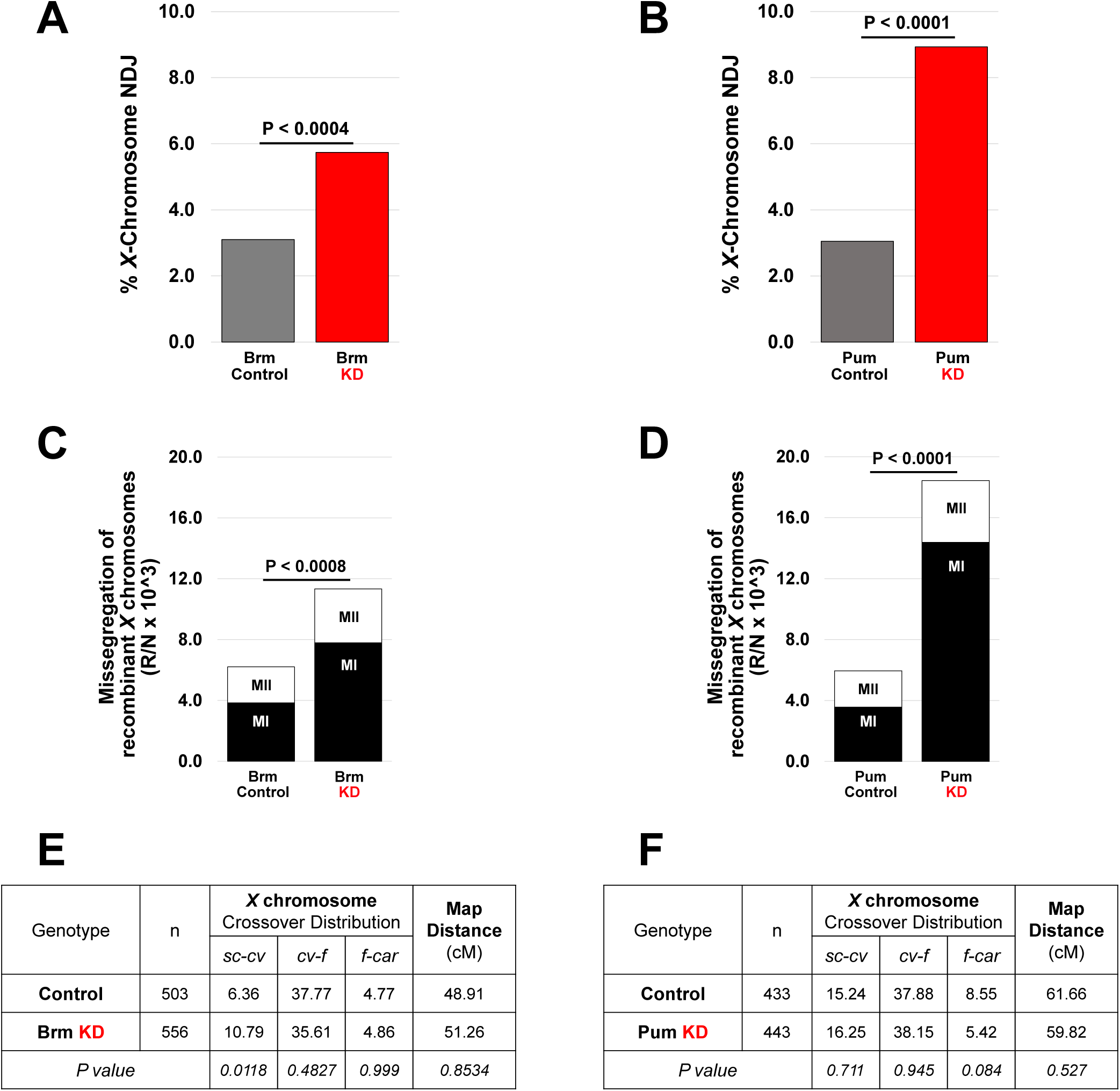
Knockdown (KD) of Brm or Pum increases missegregation of recombinant homologs without affecting crossover frequency. The matα driver and hairpins SH00130.N or SH02112.N were used to knock down Brm or Pum, respectively. **A&B.** *X*-chromosome NDJ tests for Brm KD and Pum KD (red bars) and their respective controls (gray bars) yielded results consistent with those presented in Figure 4. Diplo-*X* progeny from the NDJ tests were used for the subsequent *X*-chromosome recombinational history assay. **C&D.** The frequency at which Brm or Pum KD causes missegregation of recombinant *X* chromosomes is graphed. To calculate frequency, the number of Diplo-*X* females that received at least one recombinant chromosome (R) is divided by the total number of progeny (N) and multiplied by 1000 to simplify presentation. Inheritance of two homologs (MI segregation error) is depicted in black, and inheritance of two sisters (MII segregation error) is depicted in white. Matα-induced KD of either Brm or Pum caused a significant increase in the missegregation of recombinant homologs. P values for MI errors are shown above the bars. Differences in the frequency of MII errors between KD and control were not significant for either Brm or Pum. **E&F.** Crossovers were scored within three intervals along the *X* chromosome. The map distance (cM) is presented for each interval as well as the total number of male progeny scored for each genotype (n). Matα-induced KD of Brm or Pum does not significantly alter the total map distance on the *X* chromosome (two-tailed Fisher’s exact test).

To directly assay whether sister chromatid cohesion is prematurely disrupted when Brm or Pum are knocked down during prophase, we performed FISH on mature Drosophila oocytes (stages 13-14). Using two different probes, we scored for cohesion defects within the pericentric heterochromatin as well as a distal region on the arm of the *X* chromosome (**Fig 6A-B**). All FISH experiments utilized oocytes that were wild-type for *matrimony*. When we knocked down Brm during meiotic prophase, the percentage of oocytes with arm cohesion defects increased significantly (**Fig 6C**).

**Figure 6.**
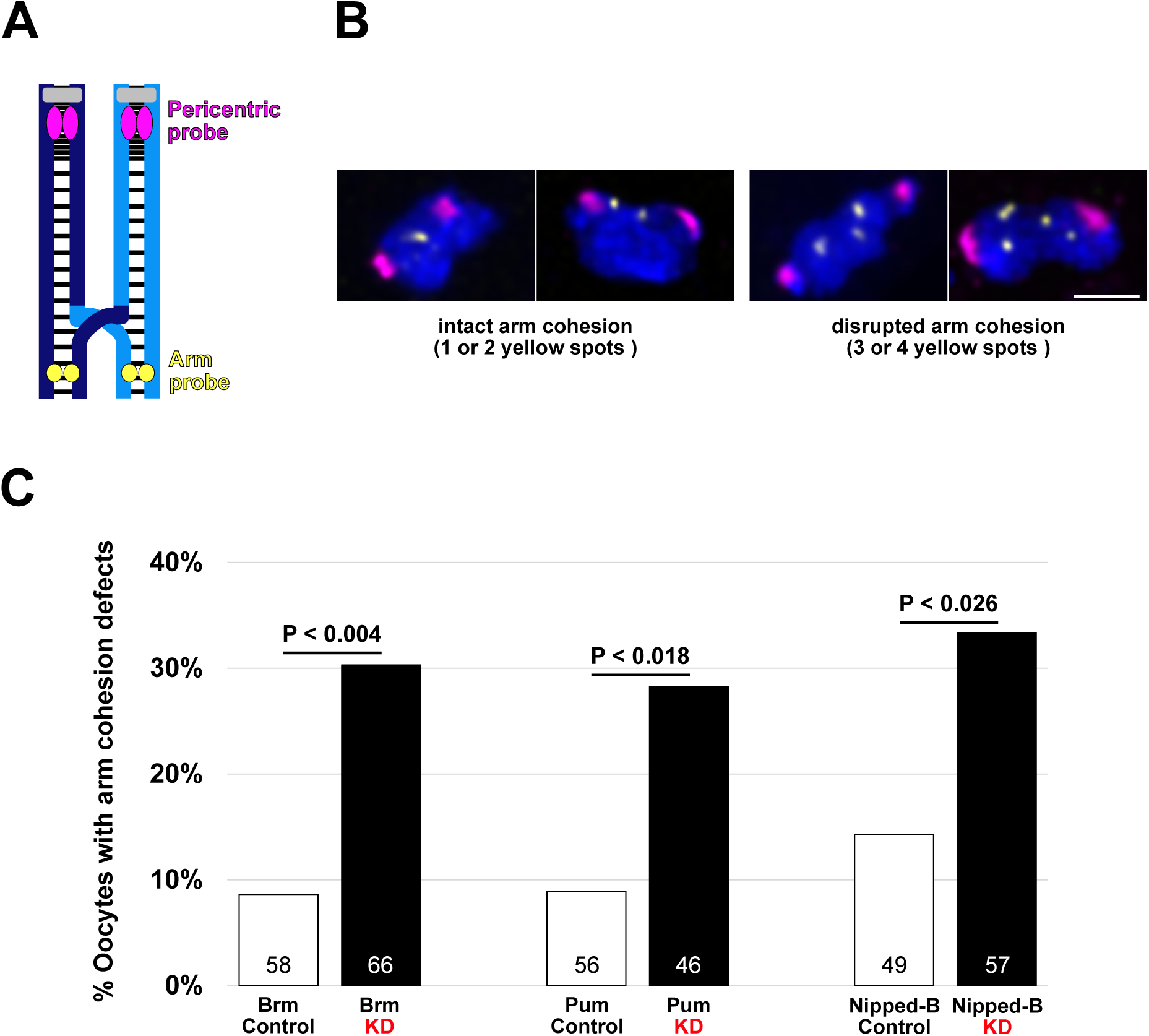
Prophase-specific positives Brm and Pum are required for cohesion maintenance during meiotic prophase. **A.** Cartoon depicts a chiasmate *X*-chromosome bivalent with intact sister chromatid cohesion. Each homolog (dark blue and light blue) is composed of two sister chromatids with sister cohesion represented by black lines and centromeres in gray. Magenta and yellow spots indicate the locations to which the FISH probes hybridize. **B.** Representative images illustrate intact or premature loss of arm cohesion in mature oocytes. Images are maximum intensity projections of deconvolved confocal Z series. Scale bar, 2µm. **C.** Quantification of arm cohesion defects in matα-Gal4 → knockdown (KD) and control (no driver) oocytes. Number of oocytes scored for each genotype is shown within each bar. No defects in pericentric cohesion were observed in any of the six genotypes tested. A two-tailed Fisher’s exact test was used to calculate P values.

Similarly, Pum KD caused a significant elevation in oocytes with premature loss of arm cohesion (**Fig 6C**). We did not detect cohesion defects in pericentric heterochromatin for either of the KD or control genotypes tested. These data indicate that Brm and Pum are required during meiotic prophase in Drosophila oocytes to maintain arm cohesion between sister chromatids, potentially by influencing Nipped-B-dependent cohesin loading during rejuvenation.

Given our previous work implicating the cohesin loader in prophase rejuvenation (HASEEB *et al*. 2023) and its functional connection with Pum and the yeast ortholog of Brm (GERBER *et al*. 2006; MUNOZ *et al*. 2019; MUNOZ *et al*. 2020), we performed FISH to directly assess the state of sister chromatid cohesion in Nipped-B KD and control oocytes. Compared to the control, the percentage of Nipped-B KD oocytes exhibiting arm cohesion defects was significantly higher (**Fig 6C**). Therefore, our FISH data confirm that Nipped-B is required during meiotic prophase for rejuvenation of arm cohesion in Drosophila oocytes. Similar to our findings for Brm and Pum, we did not observe defects in pericentric cohesion in Nipped-B KD or control oocytes. This finding aligns with our previous observation that Nipped-B localizes along the arms but not at the centromeres of oocyte chromosomes (GAUSE *et al*. 2008). We have recently reported that newly synthesized cohesin is loaded onto oocyte chromosomes during meiotic prophase and used to form new cohesive linkages (HASEEB *et al*. 2023). Together, our observations support the hypothesis that Brm and Pum activities promote Nipped-B dependent loading of cohesin onto chromosome arms during prophase rejuvenation in Drosophila oocytes.

### Additional interesting observations

Our screen also provided information about gene products required for normal reproductive biology in Drosophila females as well as the effect of hairpin insertion site on baseline NDJ in control (no KD) oocytes. Germline knockdown of several proteins severely reduced female fertility or caused complete sterility (**Table S9**). In addition, **Table S10** lists gene products for which knockdown significantly decreased NDJ compared to the control, suggesting that these proteins negatively impact the fidelity of chromosome segregation when present at normal levels in oocytes. Interestingly, we also observed that control oocytes containing an attP40 hairpin insertion exhibited higher baseline NDJ than those with an attP2 insertion (**Fig S4**). These data align with other reports that the attP40 insertion site can influence phenotypes in multiple Drosophila tissues (GROEN *et al*. 2022; VAN DER GRAAF *et al*. 2022; DUAN *et al*. 2023).

### Conclusions

Our screen, designed to identify proteins required for cohesion rejuvenation in Drosophila oocytes, uncovered 29 gene products that have a prophase-specific function required for accurate chromosome segregation. We characterized two prophase-specific positives, Brahma and Pumilio, which have functional links with the cohesin loader Nipped-B. Arm cohesion defects increase significantly when Brahma, Pumilio, or Nipped-B is knocked down during meiotic prophase, indicating that all three proteins are required for maintenance of arm cohesion in Drosophila oocytes. We propose that during prophase in Drosophila oocytes, a Brahma-containing chromatin remodeling complex recruits Nipped-B to nucleosome-free regions, facilitating the loading of new cohesin complexes onto chromosome arms. Furthermore, we posit that Pumilio-dependent stabilization of Nipped-B mRNA during meiotic prophase ensures that Nipped-B protein levels are sufficient for rejuvenation. Future analyses will better define the molecular mechanism(s) by which Brahma and Pumilio influence Nipped-B function and determine whether any of the 27 additional prophase-specific positives is required for cohesion rejuvenation. Our validated screening method also could be expanded beyond the 63 targets that we knocked down in this study.

## Author Contributions

MAH and SEB designed the experiments. MAH, ACB and EED conducted the experiments and analyzed the data. MAH and SEB wrote the manuscript.

## Data Availability Statement

Fly strains are available upon request. The authors affirm that all data necessary for confirming the conclusions of the article are present within the text, figures and tables. Supplementary Tables 1 and 2 provide details for all the fly stocks used in this study.

## Acknowledgements

We thank the Bloomington Drosophila Stock Center (NIH P40OD018537), Vienna Drosophila Resource Center (VDRC, www.vdrc.at), and the Transgenic RNAi Project (NIH R24OD030002) for fly stocks. We are grateful to the Hawley lab for providing the nanos-Gal4-VP16 stock, the Joyce lab for providing the Oligopaint arm probe and Britton Johnson for weekly fly food preparation. We also thank members of the Bickel lab (past and present) for valuable assistance and discussion. This research was funded by NIH R01GM05934 awarded to SEB.

---

**Figure S1.**
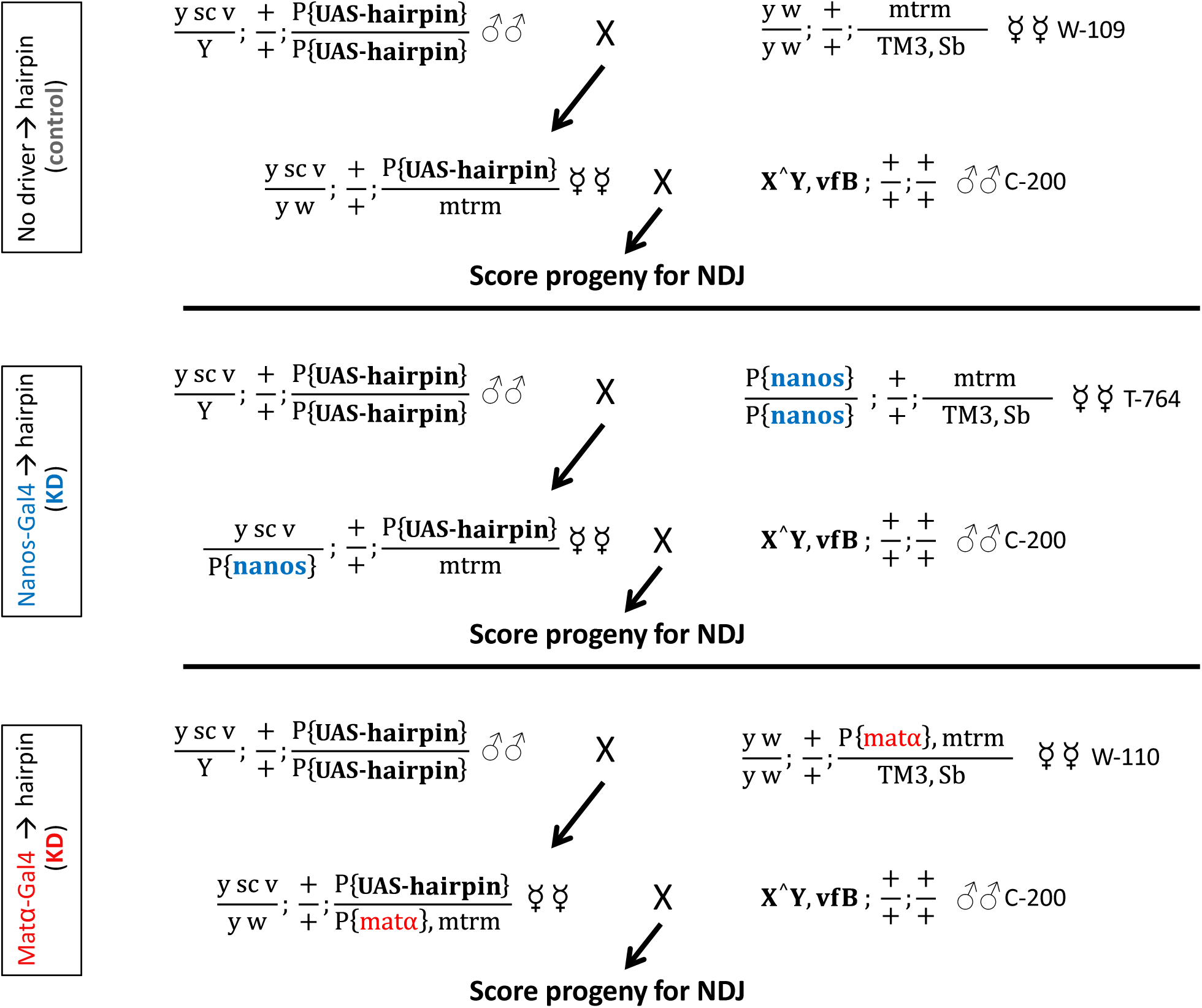
Cross schemes for NDJ screen. Crosses are shown for a hairpin stock with a *3^rd^* chromosome hairpin insertion and an *X* chromosome marked with *y sc v*. *2^nd^* chromosome hairpins were also tested, and the *X* chromosome genotype also varied between hairpin stocks. Attached *X^Y* males were crossed to Control (no driver) or KD oocytes and NDJ scored as shown in Fig 2D

**Figure S2.**
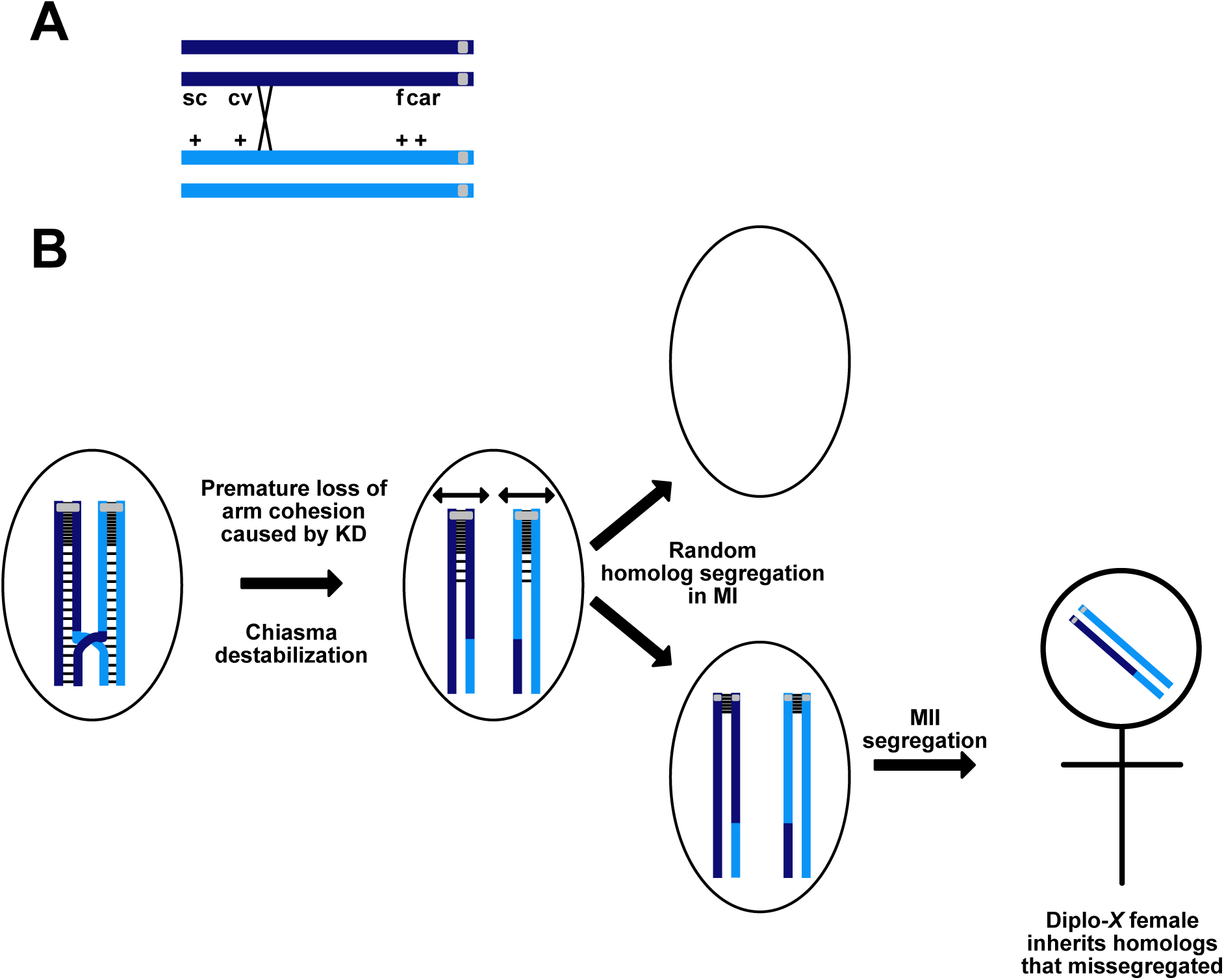
Premature loss of arm cohesion allows recombinant homologs to missegregate. **A.** In the recombinational history assay, meiotic NDJ is measured for females that are heterozygous for visible recessive markers along the *X* chromosome. Homologs, each composed of two sister chromatids, are depicted as light blue and dark blue with gray centromeres. This example illustrates an oocyte with a single crossover between *cv* and *f*. **B.** After a crossover occurs, sister chromatid cohesion (black dashed lines) keeps recombinant homologs physically associated, a prerequisite for their accurate segregation during the first meiotic division. In this example, premature loss of arm cohesion allows random segregation during MI, giving rise to a Diplo-*X* female in the NDJ assay that inherits two homologs, one of which is recombinant. By scoring the male progeny of each Diplo-*X* female for the *X*-chromosome markers, we can determine whether she inherited two homologs or sisters (based on the *car* marker), and whether either of the chromosomes is recombinant. An increase in the missegregation of recombinant homologs is consistent with premature loss of arm cohesion in KD oocytes.

**Figure S3.**
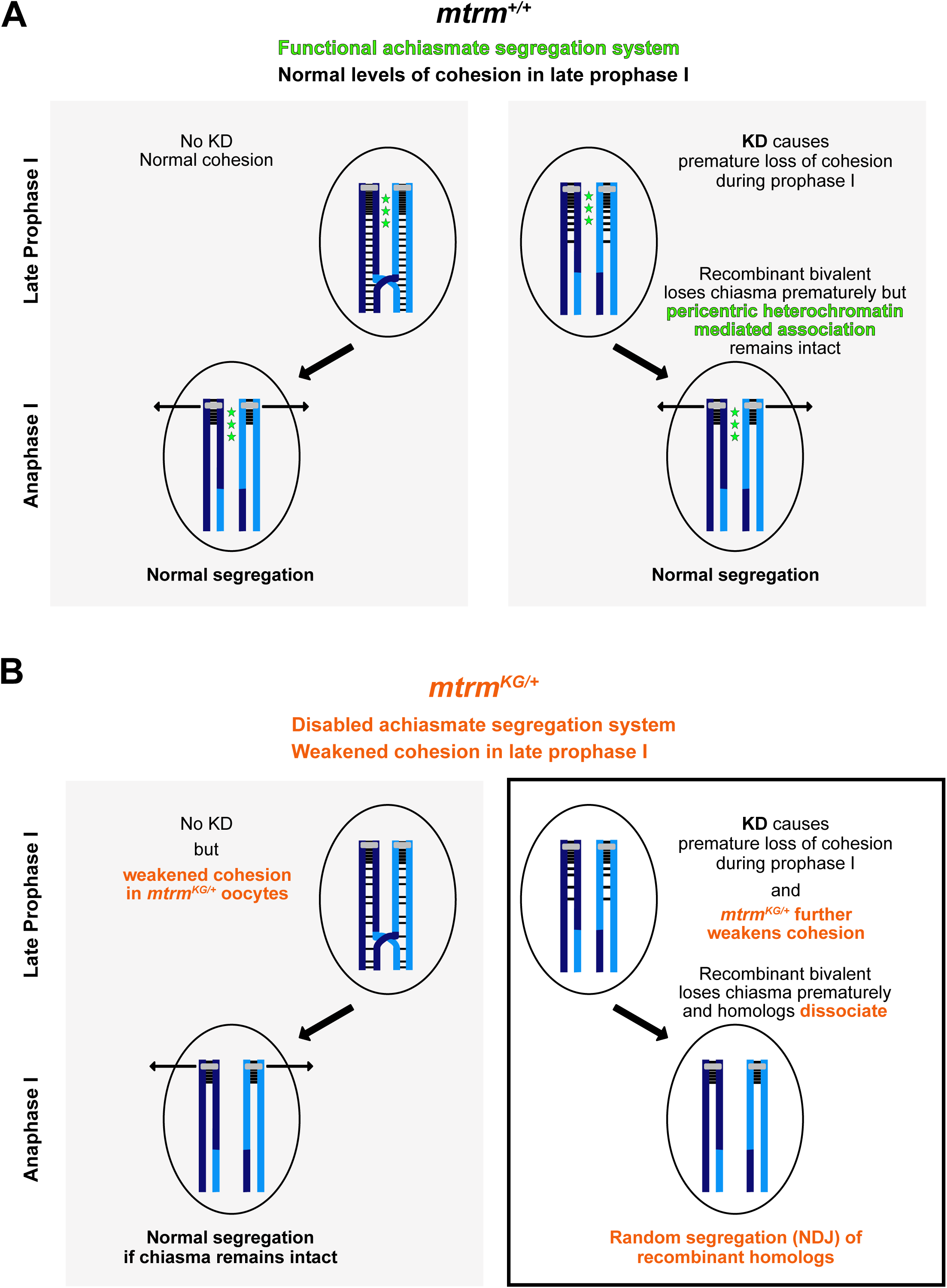
*mtrm^KG^* heterozygotes provide a sensitized genetic background to score for NDJ arising from premature loss of cohesion. **A.** In Drosophila oocytes, pericentric heterochromatin mediated association of homologs (depicted by green stars) keeps achiasmate bivalents physically associated and ensures their proper segregation (left). This same achiasmate system also promotes accurate segregation of recombinant homologs when a chiasma is destabilized due to premature loss of arm cohesion (right). Therefore, knockdown (KD) of proteins required for cohesion maintenance may not significantly increase NDJ in *mtrm^+^* oocytes. **B.** We performed our screen using oocytes that were heterozygous for the *mtrm^KG08051^* allele. The achiasmate system is disabled in *mtrm^KG/+^* oocytes (Harris et al. 2003) and cohesion is weakened (but not eliminated) due to increased Polo kinase activity in late prophase (Xiang et al. 2007; Bonner et al. 2020; Haseeb et al. 2023). In the absence of KD, weakened cohesion is still sufficient to promote accurate segregation (left). If KD further disrupts cohesion, premature dissociation of recombinant homologs leads to NDJ (right). Therefore, *mtrm^KG^* heterozygotes provide an excellent sensitized genetic background to screen for gene products that, upon knockdown, result in premature loss of cohesion.

**Figure S4.**
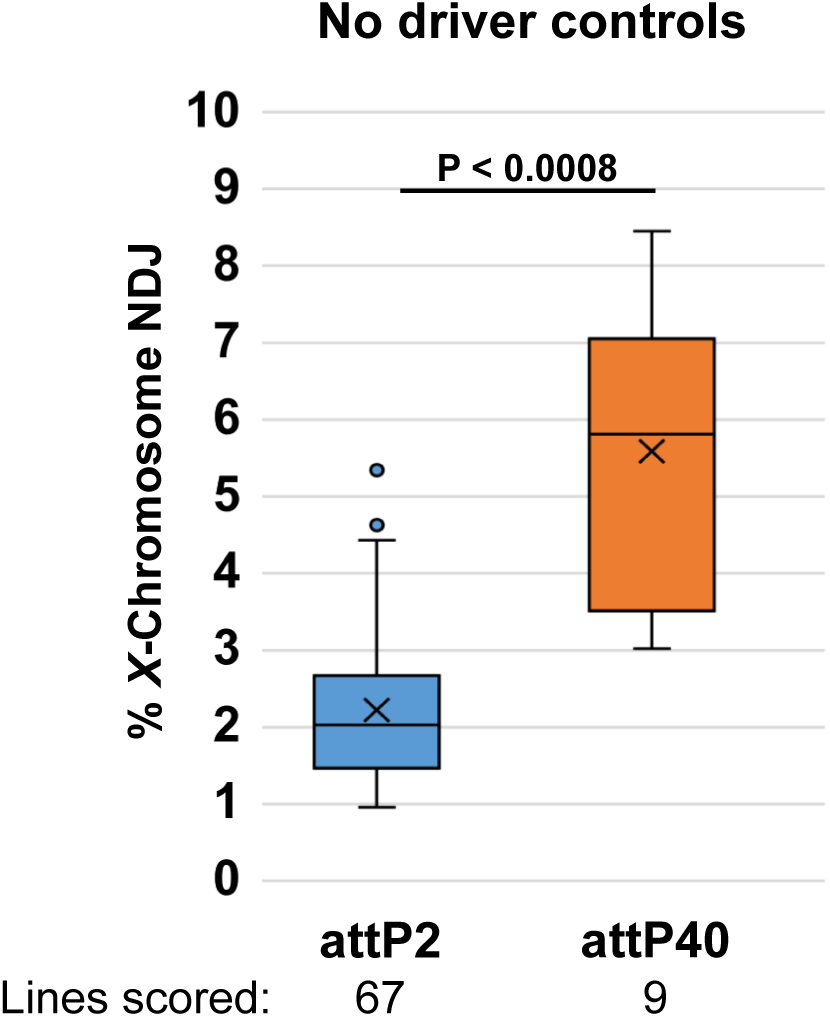
Effect of hairpin insertion site on NDJ in control oocytes. *X*-chromosome NDJ in control oocytes (no knockdown) is significantly higher for oocytes with hairpins inserted at the attP40 site on chromosome *2* than for hairpins at the attP2 site on chromosome *3*. The number of independent insertion lines tested are shown below each insertion site. An unpaired, two-tailed Student’s t-test was performed to calculate the P value.

## List of Supplementary Tables

**Table S1.** Fly stocks utilized in primary screen and subsequent assays.

| Genotype | Hairpin | Abbreviation | Source | Bickel Stock # |
| --- | --- | --- | --- | --- |
| $w^*$; + ; $P\{w^{+mC}=mata4-GAL4-VP16\}V37$ | | *mata* | BL #7063 | T-273 |
| $y^1 Df(1)w^{67c23}$ | | *y w* | | A-062 |
| $y^1$; $P\{y^{+mDint2} w^{BR.E.BR}=SUPorP\}$<br>$Exo70^{KG08051} mtrm^{KG0805} ry^{506}/$<br>$TM3,Sb^1 Ser^1$ | | | BL #14932 | M-755 |
| $y w / B^S Y$; + ; $mtrm^{KG0805}/TM3,Sb$ | | $mtrm^{KG}$ | Bickel lab derivative of M-755 | W-109 |
| $y w / B^S Y$; + ; $mtrm^{KG0805}$<br>$P\{w^{+mC}=mata4-GAL4-VP16\}V37/TM3,Sb$ | | $mtrm^{KG}, mata$ | Perkins et al. 2016 | W-110 |
| $P\{nanos-Gal4-VP16, MVD2 w^+\} y w/Y$; +<br>; $TM3,Ser,Sb/D$ | | | Hawley lab | OL-043 |
| $P\{nanos-Gal4-VP16, MVD2 w^+\} y w/B^S Y$<br>; + ; $mtrm^{KG0805}/TM3,Sb$ | | *nanos, mtrm<sup>KG</sup>* | Bickel lab derivative of M-835 & OL-043 | T-764 |
| $C(1)RM, y^2, su(w^a) w^a / X^A Y, v f B$ | | $X^A Y, Bar$ | BL #700 | C-200 |
| $w^{1118}$; $P\{w^{+mC}=UASp-Act5C.T:GFP\}2$;<br>$l(3)^{**}/TM6C, Sb^1 Tb^1$ | | | BL #7310 | B-071 |
| $y w$; $P\{w^{+mC}=UASp-Act5C.T:GFP\}2$; + | | *UASp-Actin-GFP* | Bickel Lab derivative of B-071 | A-201 |
| $y sc cv v f car/FM7a/B^S Y$; + ;<br>$mtrm^{KG0805} P\{w^{+mC}=mata4-GAL4-VP16\}V37/TM3,Sb(Ser)$ | | $y sc cv v f car$;<br>$mtrm^{KG}, mata$ | Perkins et al. 2016 | M-834 |
| $y sc cv v f car/FM7a/B^S Y$; + ;<br>$mtrm^{KG0805}/TM3,Sb(Ser)$ | | $y sc cv v f car$;<br>$mtrm^{KG}$ | Perkins et al. 2016 | M-835 |
| $y^1 sc^1 v^1$; + ; $P\{y^{+t7.7} v^{+t1.8}=TRiP$<br>$HMS00050=Brm^{V20}\}attP2$ | SH00130.N<br>Valium 20 | *TRiP Brm<sup>V20</sup>* | BL #34520 | H-209 |
| $y^1 sc^1 v^1 sev^{21}$; + ; $P\{y^{+t7.7} v^{+t1.8}=TRiP$<br>$HMS01564=Pum^{V20}\}attP2$ | SH02112.N<br>Valium 20 | *TRiP Pum<sup>V20</sup>* | BL #36676 | H-211 |
| $y^1 sc^1 v^1$; $P\{y^{+t7.7} v^{+t1.8}=$<br>$TRiP.GL00574=Nipped-B^{V22}\}attP40$; + | SH002735.N<br>Valium 22 | *TRiP Nipped-B<sup>V22</sup>* | BL #36614 | H-063 |
| $y$; + ; $P\{y^{+t7.7} v^{+t1.8}=TRiP$<br>$HMS00050=Brm^{V20}\}attP2$ | SH00130.N<br>Valium 20 | $y$; + ; *TRiP Brm<sup>V20</sup>* | Bickel lab derivative of H-209 | I-563 |
| $y$; + ; $P\{y^{+t7.7} v^{+t1.8}=TRiP$<br>$HMS01564=Pum^{V20}\}attP2$ | SH02112.N<br>Valium 20 | $y$; + ; *TRiP Pum<sup>V20</sup>* | Bickel lab derivative of H-211 | I-576 |

BL = Bloomington Drosophila Stock Center

**Table S2.** Original hairpin stocks tested in NDJ screen.

| Gene | Hairpin ID | Vector | Insertion site | Source |
| --- | --- | --- | --- | --- |
| Abl #1 | SH01558.N2 | V22 | attP2 | BL #35327 |
| Abl #2 | SH07944.N | V20 | attP40 | BL #61170 |
| AGBE | SH02397.N2 | V22 | attP2 | BL #42753 |
| AhcyL1 | SH02584.N2 | V22 | attP2 | BL #43168 |
| Baz | SH02076.N | V20 | attP2 | BL #35002 |
| Brm #1 | SH00130.N | V20 | attP2 | BL #34520 |
| Brm #2 | SH01379.N2 | V22 | attP2 | BL #35211 |
| BthD | SH03724.N2 | V22 | attP2 | BL #43267 |
| CCT3 | SH00729.N | V20 | attP2 | BL #34969 |
| Cype | SH00317.N | V20 | attP2 | BL #33878 |
| Dhc64C | SH02710.N | V20 | attP2 | BL #36698 |
| Diap1 | SH00680.N | V20 | attP2 | BL #33597 |
| EIF5 | SH00345.N | V20 | attP2 | BL #34841 |
| Fmr1 #1 | SH00354.N | V20 | attP2 | BL #34944 |
| Fmr1 #2 | SH01274.N2 | V22 | attP2 | BL #35200 |
| Fs(1)k10 | SH00087.N | V20 | attP2 | BL #33630 |
| Grp | SH01903.N | V20 | attP2 | BL #36685 |
| Gwl | SH00920.N | V20 | attP2 | BL #34525 |
| Hang | SH02893.N2 | V22 | attP40 | BL #41870 |
| HEM | SH04538.N | V20 | attP2 | BL #41688 |
| Hip14 | SH02106.N | V20 | attP2 | BL #35012 |
| ifc | SH00391.N | V20 | attP2 | BL #32514 |
| Imp | SH01803.N | V20 | attP2 | BL #34977 |
| Iswi | SH00393.N | V20 | attP2 | BL #32845 |
| Lsd-2 | SH00412.N | V20 | attP2 | BL #32846 |
| Mamo | SH07656.N | V20 | attP2 | BL #60111 |
| Mbc | SH05347.N | V20 | attP2 | BL #36658 |
| mCherry | SH02163.N | V20 | attP2 | BL #35785 |
| Mps1 #1 | SH01602.N | V20 | attP2 | BL #36658 |
| Mps1 #2 | shRNA-330259 | Walium 20 | attP40 | VDRC #330259 |
| Ord | SH02738.N2 | V22 | attP2 | BL #36633 |
| pAbp | unavailable | V20 | attP2 | BL #67979 |
| Park | SH03863.N | V20 | attP2 | BL #38333 |
| Plc21C | SH00900.N | V20 | attP2 | BL #33719 |
| Pum #1 | SH02803.N | V20 | attP40 | BL #38241 |
| Pum #2 | SH02112.N | V20 | attP2 | BL #36676 |
| Punch | SH03835.N | V20 | attP2 | BL #55397 |
| Raf | SH03151.N | V20 | attP2 | BL #55679 |
| Rap1 | SH02250.N | V20 | attP2 | BL #35047 |
| Ras85D | SH01307.N | V20 | attP2 | BL #34619 |
| Rbfox1 | SH00782.N | V20 | attP2 | BL #32476 |
| RhoGAP1A | SH00517.N | V20 | attP2 | BL #33390 |
| RhoGAP92B | SH00519.N | V20 | attP2 | BL #33391 |
| RYBP | SH00691.N | V20 | attP2 | BL #33974 |
| Scf | SH01884.N | V20 | attP2 | BL #34331 |
| Septin 4 | SH02657.N2 | V22 | attP2 | BL #41859 |
| SERCA | SH004955.N | V20 | attP2 | BL #44581 |
| Singed | SH04535.N | V22 | attP2 | BL #42615 |
| SmB | SH06421.N | V20 | attP2 | BL #64594 |
| Smc1 | SH01950.N2 | V22 | attP2 | BL #36598 |
| Smc2 | SH00559.N | V20 | attP2 | BL #32369 |
| Smc3 | SH03166.N | V20 | attP40 | BL #50899 |
| Smc4 | SH00364.N | V20 | attP2 | BL #32350 |
| Socs44A | SH04771.N | V20 | attP2 | BL #42830 |
| Sunn | SH020-F10 | V20 | attP40 | BL #52969 |
| Tao | SH01902.N | V20 | attP2 | BL #34881 |
| Thiolase | SH00737.Nb | V20 | attP2 | BL #34546 |
| Tos | SH00792.N | V20 | attP2 | BL #33937 |
| Tre1 | SH00895.N | V20 | attP2 | BL #34956 |
| CG2941 | SH02890.N2 | V22 | attP40 | BL #41867 |
| CG4294 | SH00243.N | V20 | attP2 | BL #33357 |
| CG5292 #1 | SH00947.N | V20 | attP2 | BL #32499 |
| CG5292 #2 | SH00948.N | V20 | attP2 | BL #32500 |
| CG6418 | SH00920.N | V20 | attP2 | BL #32441 |
| CG6805 #1 | SH00873.N | V20 | attP2 | BL #32380 |
| CG6805 #2 | SH00872.N | V20 | attP2 | BL #34615 |
| CG7115 | SH08133.N | V20 | attP2 | BL #60015 |
| CG9925 | SH01506.Nb | V20 | attP2 | BL #35811 |
| CG10082 #1 | SH00874.N | V20 | attP2 | BL #32431 |
| CG10082 #2 | SH00875.N | V20 | attP2 | BL #33717 |
| CG10924 | SH00155.N | V20 | attP2 | BL #36915 |
| CG12084 | SH01649.N | V20 | attP2 | BL #34553 |
| CG14712 | SH01504.N2 | V22 | attP2 | BL #35622 |
| CG17658 | SH01091.N2 | V22 | attP40 | BL #36872 |
| CG18446 | SH00207.N | V20 | attP2 | BL #33735 |
| CG42232 | SH07828.N2 | V20 | attP40 | BL #64604 |

BL = Bloomington Drosophila Stock Center
VDRC = Vienna Drosophila Resource Center
V20 and V22 are TRiP vectors Valium 20 and Valium 22 respectively

**Table S3.** Gene products for which nanos-induced knockdown causes significantly higher NDJ than the matα driver.

| Gene name (hairpin ID)<br>*Vector, insertion site* | % X-chromosome NDJ<br>(Fertility) |  |  | P value |  |  |
| --- | --- | --- | --- | --- | --- | --- |
| | Control | Nanos KD | $\text{Mat}\alpha$ KD | Nanos<br>&<br>Control | $\text{Mat}\alpha$<br>&<br>Control | Nanos<br>&<br>$\text{Mat}\alpha$ |
| **Smc1** (SH01950.N2)<br>*V22, attP2* | 1.57<br>(11.1) | *35.24<br>(16.4) | *6.08<br>(16.8) | <0.0001 | <0.0001 | <0.0001 |
| **Smc3** (SH03166.N)<br>*V20, attP40* | 7.80<br>(15.1) | *33.69<br>(13.5) | *18.33<br>(13.8) | <0.0001 | <0.0001 | <0.0001 |
| **Ord** (SH02738.N2)<br>*V22, attP2* | 2.35<br>(19.5) | *57.19<br>(5.30) | *5.62<br>(17.7) | <0.0001 | 0.0017 | <0.0001 |
| **Sunn** (SH020-F10)<br>*V20, attP40* | 3.02<br>(14.7) | *47.62<br>(4.60) | *9.29<br>(16.9) | <0.0001 | <0.0001 | <0.0001 |
| **Rbfox1** (SH00782.N)<br>*V20, attP2* | 1.69<br>(13.2) | *34.78<br>(13.6) | *6.47<br>(13.8) | <0.0001 | <0.0001 | <0.0001 |
| **Mamo** (SH07656.N)<br>*V20, attP2* | 1.73<br>(11.5) | *15.37<br>(14.3) | *6.67<br>(12.5) | <0.0001 | <0.0001 | <0.0001 |
| **Negative Control:** |  |  |  |  |  |  |
| **mCherry** (SH02163.N)<br>*V20, attP2* | 2.30<br>(15.1) | 2.18<br>(18.2) | 3.31<br>(18.6) | 0.88 | 0.26 | 0.18 |

Fertility values shown in ( ) indicate the number of progeny per female in the NDJ assay. Asterisk indicates a significant difference in NDJ compared to the control ( $P < 0.05$ ). V20 and V22 are VALIUM 20 and VALIUM 22 vectors respectively.

**Table S4.** Elevated NDJ is comparable for the two drivers.

| Gene name (hairpin ID)<br>*Vector, insertion site* | % X-chromosome NDJ<br>*(Fertility)* |  |  | P value |  |  |
| --- | --- | --- | --- | --- | --- | --- |
| | Control | Nanos KD | Matα KD | Nanos<br>&<br>Control | Matα<br>&<br>Control | Nanos<br>&<br>Matα |
| **Fmr1 #1** (SH00354.N)<br>*V20, attP2* | 1.56<br>(14.3) | *8.77<br>(18.0) | *10.26<br>(9.00) | <0.0001 | <0.0001 | 0.44 |
| **Fmr1 #2** (SH01274.N2)<br>*V22, attP2* | 3.08<br>(12.0) | *6.45<br>(10.1) | *6.31<br>(9.20) | 0.021 | 0.031 | 0.935 |
| **BthD** (SH03724.N2)<br>*V22, attP2* | 1.99<br>(7.50) | *6.74<br>(10.4) | *10.18<br>(13.1) | 0.0013 | <0.0001 | 0.057 |
| **Ras85D** (SH01307.N)<br>*V20, attP2* | 1.36<br>(20.1) | *9.28<br>(20.3) | *6.96<br>(17.7) | <0.0001 | <0.0001 | 0.10 |
| **Scf** (SH01884.N)<br>*V20, attP2* | 3.96<br>(17.3) | *7.52<br>(17.9) | *8.54<br>(17.7) | 0.0039 | <0.0004 | 0.48 |
| **Punch** (SH03835.N)<br>*V20, attP2* | 1.30<br>(15.3) | *4.59<br>(16.0) | *8.18<br>(18.8) | <0.0006 | <0.0001 | 0.0057 |
| **AGBE** (SH02397.N2)<br>*V22, attP2* | 2.40<br>(11.3) | *5.66<br>(12.0) | *6.29<br>(10.0) | 0.011 | 0.0058 | 0.70 |
| **Raf** (SH03151.N)<br>*V20, attP2* | 2.67<br>(11.1) | *5.44<br>(13.9) | *5.90<br>(13.2) | 0.025 | 0.012 | 0.74 |
| **Iswi** (SH00393.N)<br>*V20, attP2* | 1.09<br>(11.4) | *5.31<br>(10.1) | *4.58<br>(17.6) | <0.0006 | <0.0002 | 0.59 |
| **CG4294** (SH00243.N)<br>*V20, attP2* | 1.39<br>(10.7) | *3.61<br>(12.9) | *4.78<br>(14.3) | 0.03 | 0.0014 | 0.33 |
| **CG9925** (SH01506.Nb)<br>*V20, attP2* | 0.96<br>(13.0) | *2.91<br>(15.2) | *3.69<br>(17.3) | 0.015 | 0.0011 | 0.43 |

*Fertility values* shown in ( ) indicate the number of progeny per female in the NDJ assay. Asterisk indicates a significant difference in NDJ compared to the control (P < 0.05). V20 and V22 are VALIUM 20 and VALIUM 22 vectors respectively.

**Table S5.** Hairpins for which only the nanos driver significantly increases NDJ.

| Gene name (hairpin ID)<br>*Vector, insertion site* | % X-chromosome NDJ<br>*(Fertility)* |  |  | P value |  |  |
| --- | --- | --- | --- | --- | --- | --- |
| | Control | Nanos KD | Mataα KD | Nanos<br>&<br>Control | Mataα<br>&<br>Control | Nanos<br>&<br>Mataα |
| **HEM** (SH04538.N)<br>*V20, attP2* | 2.38<br>(10.4) | *5.48<br>(14.7) | 3.45<br>(13.5) | 0.0099 | 0.32 | 0.098 |
| **Cype** (SH00317.N)<br>*V20, attP2* | 1.46<br>(17.0) | *4.22<br>(20.3) | 1.46<br>(17.0) | 0.0010 | 0.99 | 0.0010 |
| **Baz** (SH02076.N)<br>*V20, attP2* | 2.41<br>(20.5) | *4.21<br>(18.6) | 4.10<br>(20.3) | 0.048 | 0.055 | 0.91 |

*Fertility values* shown in ( ) indicate the number of progeny per female in the NDJ assay. Asterisk indicates a significant difference in NDJ compared to the control (P < 0.05). V20 and V22 are VALIUM 20 and VALIUM 22 vectors respectively.

**Table S6.**
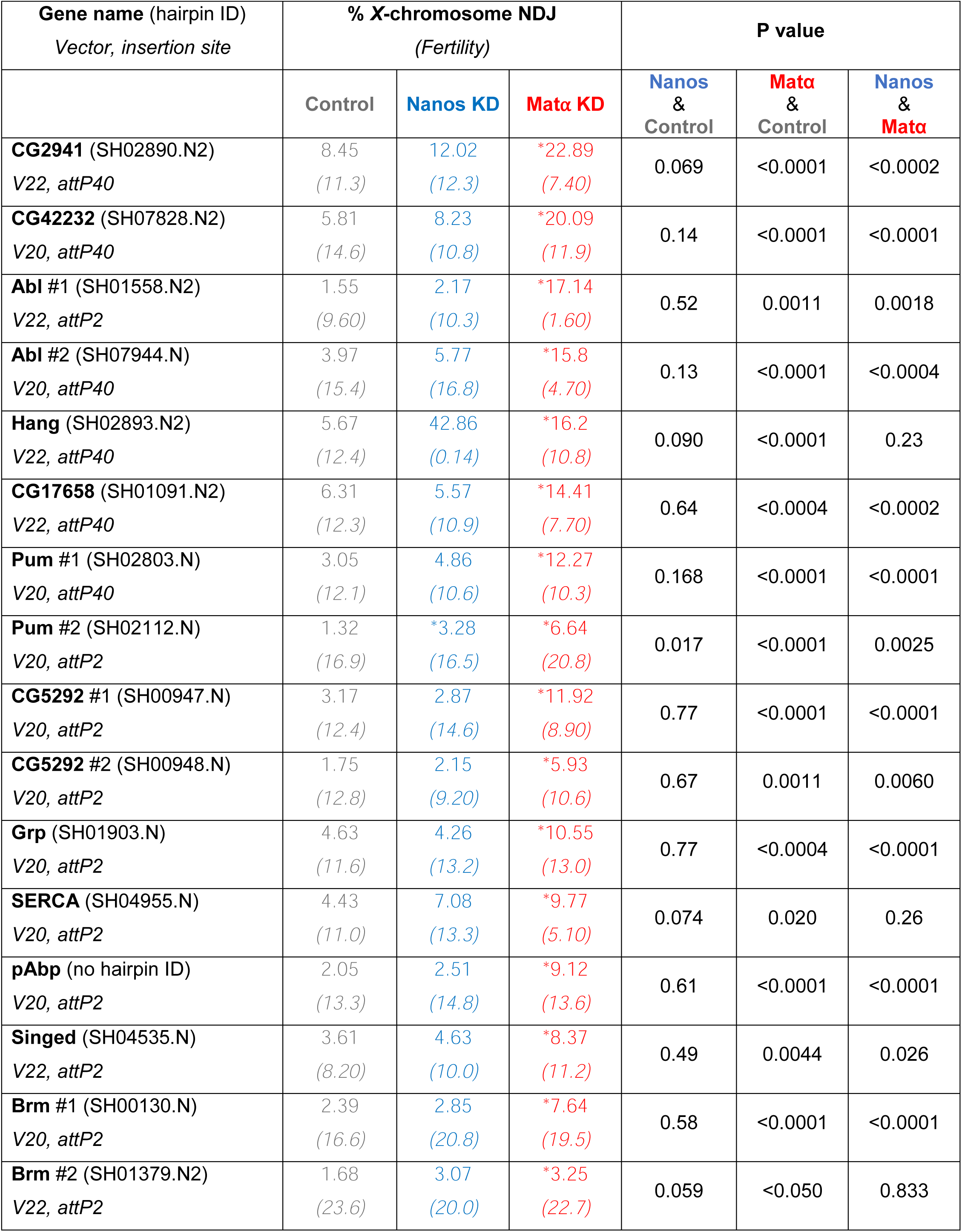
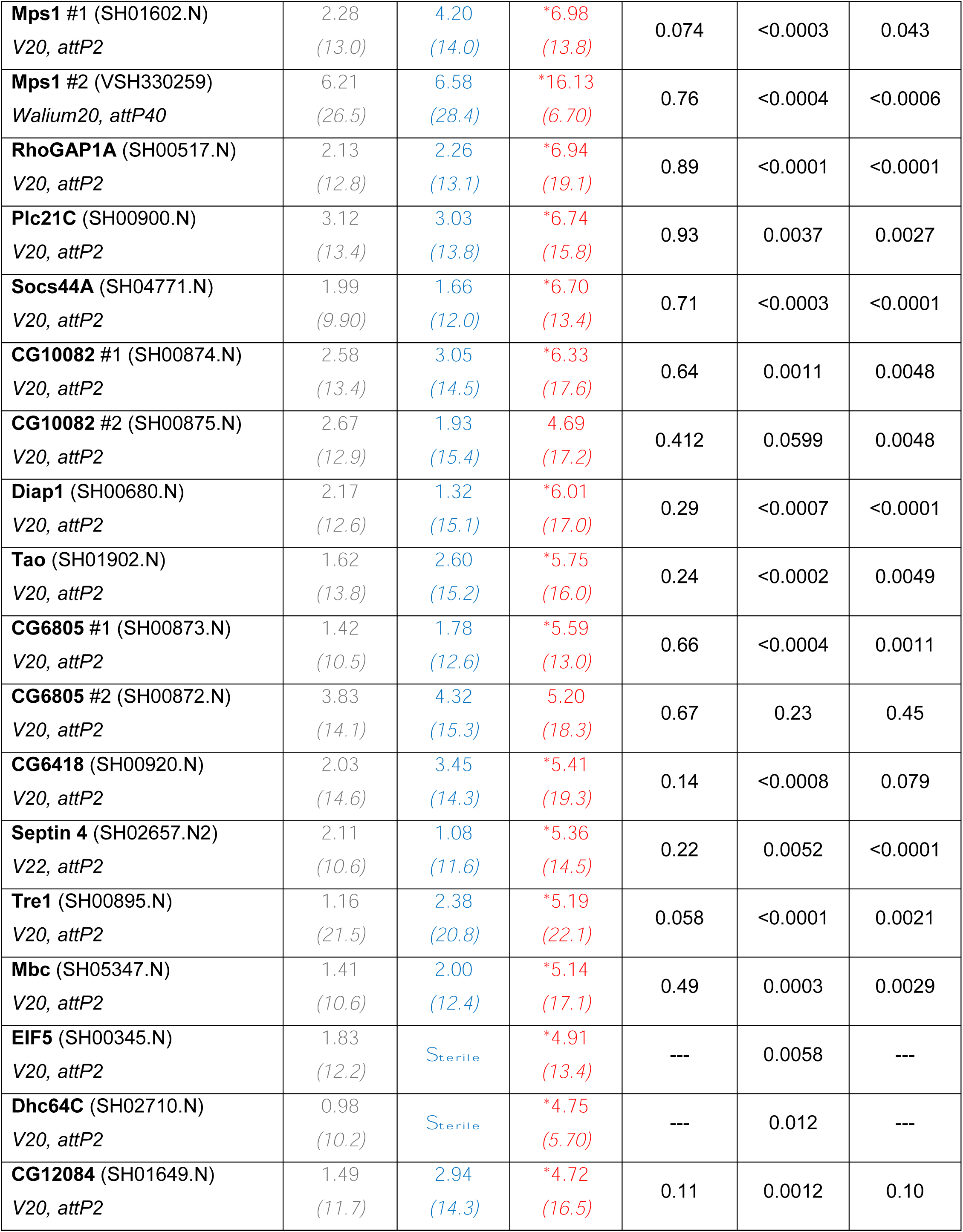
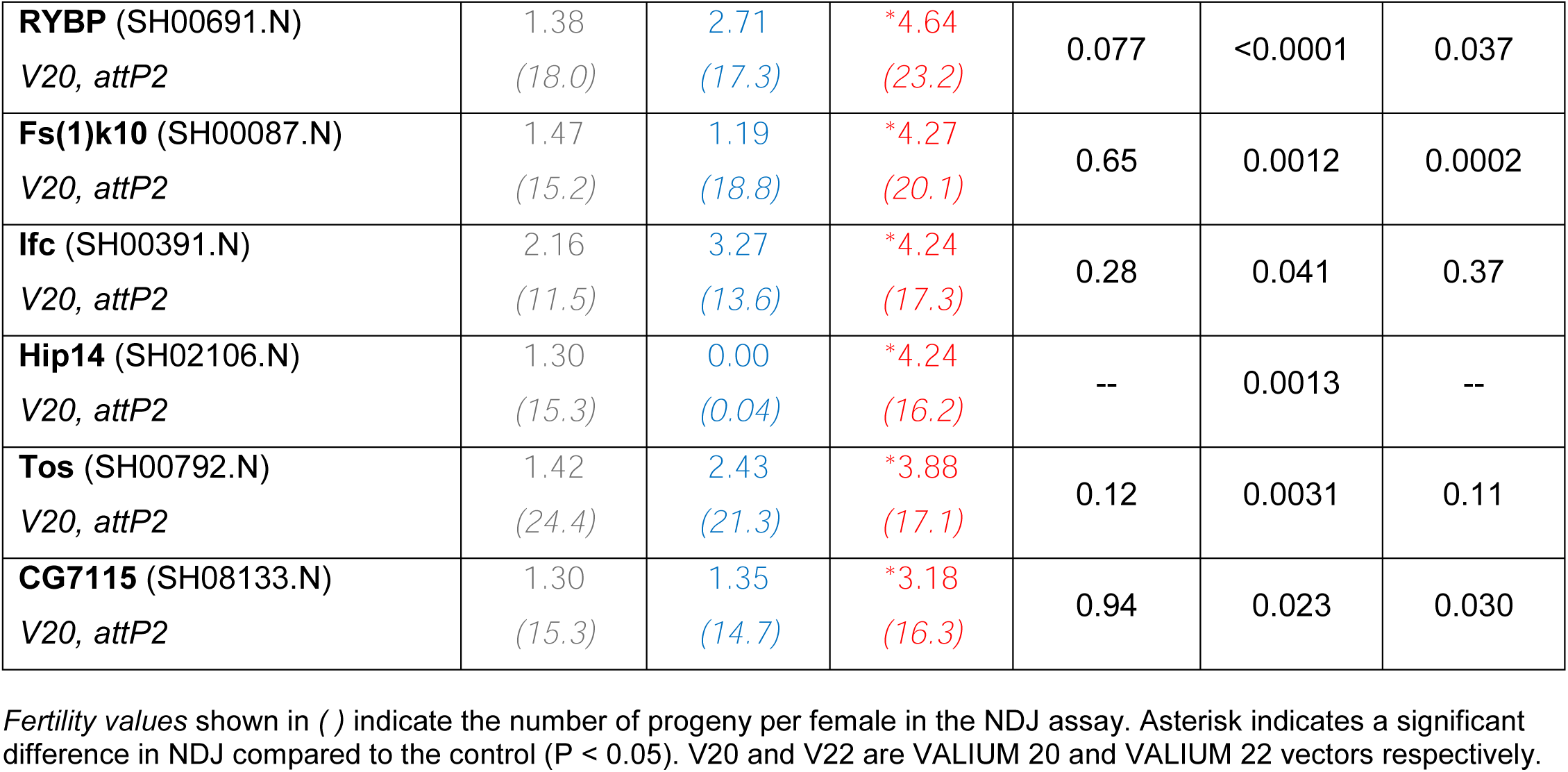
NDJ and fertility data for prophase-specific positives.

**Table S7.** Mat⍺ negatives.

| Gene name (hairpin ID)<br>*Vector, insertion site* | % X-chromosome NDJ<br>*(Fertility)* |  |  | P value |  |  |
| --- | --- | --- | --- | --- | --- | --- |
| | Control | Nanos KD | Matα KD | Nanos<br>&<br>Control | Matα<br>&<br>Control | Nanos<br>&<br>Matα |
| **Baz** (SH02076.N)<br>*V20, attP2* | 2.41<br>(20.5) | *4.21<br>(18.6) | 4.10<br>(20.3) | 0.048 | 0.055 | 0.91 |
| **CG10924** (SH00155.N)<br>*V20, attP2* | 2.24<br>(15.4) | 2.31<br>(16.1) | 2.61<br>(18.9) | 0.94 | 0.66 | 0.71 |
| **CG18446** (SH00207.N)<br>*V20, attP2* | 1.85<br>(10.7) | 2.11<br>(11.7) | 2.44<br>(12.2) | 0.78 | 0.54 | 0.73 |
| **Cype** (SH00317.N)<br>*V20, attP2* | 1.46<br>(17.0) | *4.22<br>(20.3) | 1.46<br>(17.0) | 0.0010 | 0.99 | 0.0010 |
| **HEM** (SH04538.N)<br>*V20, attP2* | 2.38<br>(10.4) | *5.48<br>(14.7) | 3.45<br>(13.5) | 0.0099 | 0.32 | 0.098 |
| **Lsd-2** (SH00412.N)<br>*V20, attP2* | 1.66<br>(15.9) | *0.17<br>(15.0) | 2.47<br>(15.0) | 0.0095 | 0.33 | <0.0005 |
| **Park** (SH03863.N)<br>*V20, attP2* | 5.34<br>(10.9) | *2.01<br>(12.3) | 8.32<br>(12.1) | 0.0077 | 0.072 | <0.0001 |
| **RhoGAP92B** (SH00519.N)<br>*V20, attP2* | 4.35<br>(11.9) | 2.86<br>(12.9) | 3.46<br>(12.8) | 0.24 | 0.50 | 0.58 |
| **SmB** (SH06421.N)<br>*V20, attP2* | 2.81<br>(14.0) | Sterile | 2.62<br>(12.2) | -- | 0.85 | -- |
| **Smc4** (SH00364.N)<br>*V20, attP2* | 2.75<br>(13.5) | Sterile | 3.44<br>(6.40) | -- | 0.61 | -- |
| **Thiolase** (SH00737.Nb)<br>*V20, attP2* | 1.32<br>(13.2) | 0.64<br>(15.6) | 1.87<br>(15.9) | 0.25 | 0.45 | 0.048 |

*Fertility values* shown in ( ) indicate the number of progeny per female in the NDJ assay. Asterisk indicates a significant difference in NDJ compared to the control (P < 0.05). V20 and V22 are VALIUM 20 and VALIUM 22 vectors respectively.

**Table S8.** Best described cellular function for each prophase-specific positive.

| **Gene name** | **Cellular function** |
| --- | --- |
| CG10082 | Predicted kinase activity |
| Abl | Kinase activity |
| Grp | Kinase activity |
| Mps1 | Kinase activity |
| Tao | Kinase activity |
| CG6805 | Predicted phosphatase activity |
| CG7115 | Predicted phosphatase activity |
| pAbp | mRNA binding |
| Pum | mRNA binding |
| Brm | Chromatin remodeling |
| CG12084 | Predicted ubiquitination |
| Diap1 | Ubiquitination |
| RYBP | Ubiquitination |
| Socs44A | Ubiquitination |
| Fs(1)k10 | Gametogenesis |
| Ifc | Gametogenesis |
| Mbc | GTPase activity |
| RhoGAP1A | GTPase activity |
| Septin 4 | GTPase activity |
| Plc21C | Calcium ion homeostasis |
| SERCA | Calcium ion homeostasis |
| CG5292 | Miscellaneous - Predicted tRNA deaminase activity |
| CG6418 | Miscellaneous - Predicted RNA helicase |
| Singed | Miscellaneous - Actin binding |
| Tre1 | Miscellaneous - G Protein coupled receptor activity |
| Tos | Miscellaneous - DNA exonuclease |
| CG2941 | Unknown |
| CG42232 | Unknown |
| CG17658 | Unknown |

**Table S9.** Hairpins for which knockdown decreases fertility at least three-fold.

| Decreased fertility with mataα driver | | |
| --- | --- | --- |
| Gene name | Control Fertility | Mataα Fertility |
| CG14712 | 14.3 | 0.4 |
| CCT3 | 15.4 | 1.1 |
| Gwl | 17.5 | 1.7 |
| Smc2 | 10.7 | 1.7 |
| Imp | 15.9 | 2.6 |
| Rap1 | 17.7 | 3.0 |
| Abl hairpin#1 | 9.6 | 1.6 |
| Abl hairpin#2 | 15.4 | 4.7 |
| Mps1 hairpin#2 | 26.5 | 6.7 |
| Decreased fertility with nanos driver |  |  |
| Gene name | Control Fertility | Nanos Fertility |
| EIF5 | 12.2 | Sterile |
| Dhc64C | 10.2 | Sterile |
| CCT3 | 15.4 | Sterile |
| SmB | 14.0 | Sterile |
| Smc2 | 10.7 | Sterile |
| Smc4 | 13.5 | Sterile |
| CG14712 | 14.3 | 0.04 |
| Hip14 | 15.3 | 0.04 |
| Gwl | 17.5 | 0.10 |
| Hang | 12.4 | 0.14 |
| AhcyL1 | 12.2 | 0.80 |
| Sunn | 14.7 | 4.60 |
| Ord | 19.5 | 5.30 |

Fertility values indicate number of progeny per female in the NDJ assay.

**Table S10.** Hairpins for which knockdown reduces NDJ.

| Gene name (hairpin ID)<br>*Vector, insertion site* | % X-chromosome NDJ<br>*(Fertility)* |  |  | P value |  |  |
| --- | --- | --- | --- | --- | --- | --- |
| | Control | Nanos KD | Mataα KD | Nanos<br>&<br>Control | Mataα<br>&<br>Control | Nanos<br>&<br>Mataα |
| **Lsd-2** (SH00412.N)<br>*V20, attP2* | 1.66<br>(15.9) | *0.17<br>(15.0) | 2.47<br>(15.0) | 0.0095 | 0.33 | 0.00044 |
| **Park** (SH03863.N)<br>*V20, attP2* | 5.34<br>(10.9) | *2.01<br>(12.3) | 8.32<br>(12.1) | 0.0077 | 0.072 | <0.0001 |
| **AhcyL1** (SH02584.N2)<br>*V22, attP2* | 3.82<br>(12.2) | 8.57<br>(0.80) | *1.36<br>(5.50) | 0.33 | 0.036 | 0.14 |

*Fertility values* shown in ( ) indicate the number of progeny per female in the NDJ assay. Asterisk indicates a significant difference in NDJ compared to the control (P < 0.05). V20 and V22 are VALIUM 20 and VALIUM 22 vectors respectively.

